# Short heat shock has a long-term effect on mesenchymal stem cells’ transcriptome

**DOI:** 10.1101/2022.11.19.517224

**Authors:** Ivana Ribarski-Chorev, Gisele Schudy, Carmit Strauss, Sharon Schlesinger

**Affiliations:** The Robert H. Smith Faculty of Agriculture, Food and Environment, The Hebrew University of Jerusalem, Israel

**Keywords:** Heat shock, bovine, mesenchymal stem cells, transcriptome, stress, bivalent genes, umbilical cord.

## Abstract

The harmful consequences of heat stress (HS) on physiology are well documented, but the molecular aspects of changing thermal conditions are poorly understood. Therefore, a better understanding of the effects of this stress on the morphology, phenotype, proliferative capacity, and fate decision of MSCs is required. Our thorough characterization of MSCs’ transcriptome showed a major effect of HS on the transcriptional landscape. Specifically, examining the effect after three days of moderate HS shows changes in many cell processes, such as immune response, cell cycle, and differentiation. Surprisingly, we detected a long-term effect on cell identity even after short stress, possibly through the activation of bivalent genes related to cell lineage decisions. Finally, comparing the differentially expressed genes following short HS with their transcriptional state after three days of recovery, we find transient upregulation of many members of the MLL family and other histone modifiers; a finding which offers a potential mechanistic account for the stable bivalent genes activation. This could be used to predict and modify the long-term effect of HS on cell identity.

**Summary blurb:** Heat shock alters mesenchymal stem cells’ transcriptional programs, resulting in stable activation of lineage commitment genes, thus explaining the shift in the identity and fate of the cells.

**Graphical abstract:** 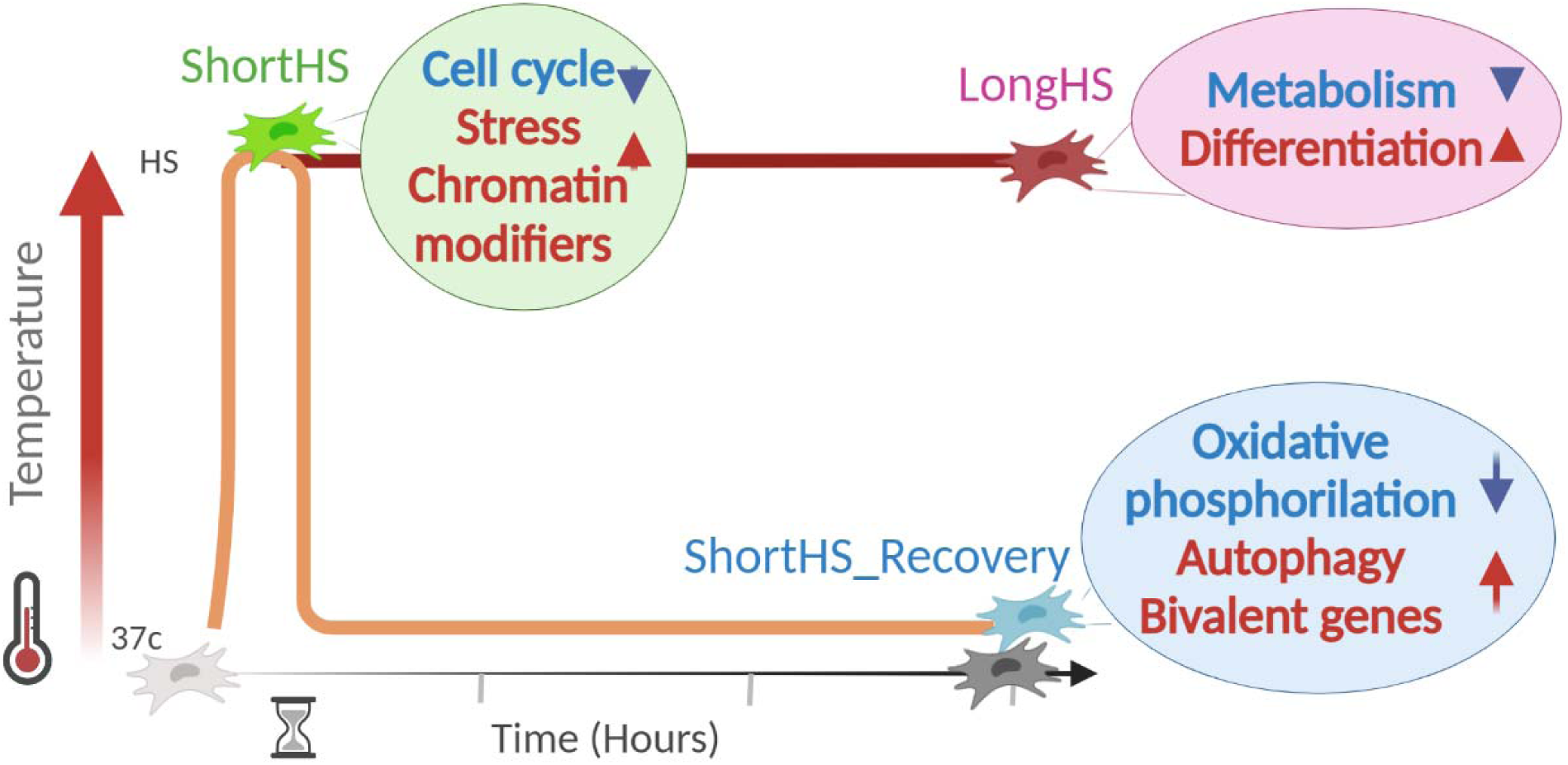

## Introduction

Environmental stressors harm human and animal health around the world (Peters et al., 2021; Richter et al., 2010). Heat exposure in particular is an increasing problem as climate change progresses; exposures to extreme temperatures vastly affect the organism as a whole and on the cellular level. Adverse effects of heat stress in humans include heat stroke, systemic inflammation, nervous system impairment, adverse birth outcomes, global DNA methylation, and telomere shortening (Peters et al., 2021). In animals, heat stress was found to have detrimental effects on fertility and well-being (Agarwal et al., 2014; de Barros & Paula-Lopes, 2018; Kitagawa et al., 2004). In cattle, the frequency of many chronic inflammation-related diseases is elevated during hot periods (Chen et al., 2018; Olde Riekerink et al., 2007), resulting in reduced animal welfare and significant economic losses to the dairy industry (Key et al., 2014). In-utero heat stress in cattle was recently found to reduce placental weight, blood flow, birth weight, and innate and cellular immunity (Dado-Senn et al., 2020). Moreover, heat stress has a long-term effect on oocyte developmental capacity and follicular steroidogenesis in cattle and other mammals (Kawano et al., 2022; Roth, 2017). It has been suggested that alterations of the cellular epigenetic landscape are responsible for these persisting effects (de Barros & Paula-Lopes, 2018; Sharma & Bhonde, 2020; Xue & Acar, 2018).

On a cellular level, thermal stress is associated with oxidative stress, endoplasmic reticulum (ER) stress, and neurochemical stress, and impairs protein folding, cell cycle, and mitochondrial function (Agarwal et al., 2014; Chen et al., 2018; Kitagawa et al., 2004; Shimoni et al., 2020). On the transcriptional level, cells exposed to HS generally activate a cascade of heat shock proteins and factors which in turn regulate downstream pathways that protect the stressed cells. The effect of external temperature has been examined in various animals and tissues such as bovine PBMCs (Garner et al., 2020), mammary tissues (Collier et al., 2006; Yue et al., 2020) and granulosa cells (Sammad et al., 2022). Another study exposed rats to heat stress and examined the transcriptional changes in 3 different tissues (Dou et al., 2021). Although many key genes and pathways that respond to acute thermal stress were identified in these studies, only a few genes were common across tissues, suggesting a high level of tissue specificity in the transcriptional response to thermal stress.

Heat stress effects on adult stem cells are of special interest. These cells are the longest-living proliferative cells in multicellular organisms (Schultz & Sinclair, 2016); hence they have an increased risk of accumulating genetic and metabolic damage. Extrinsic factors including environmental stress can enhance this damage accumulation, possibly leading to the functional decline of the stem cells (Ermolaeva et al., 2018). Mesenchymal stromal cells (also called mesenchymal stem cells, MSCs) are a heterogeneous group of non-hematopoietic multipotent stem cells that assist in the preservation of homeostasis in many organs and tissues (Pittenger et al., 1999). Additionally, physiological MSC stores are essential for the regeneration of many tissues in the body, acting through common signaling pathways such as Wnt, BMP, and Notch (Naaldijk et al., 2015). Cultured MSCs are most frequently derived from adult tissue sources such as bone marrow and adipose tissue or from birth-associated tissue such as placental tissue, amniotic membranes, and umbilical cord (Hass et al., 2011; Nowakowski et al., 2016). When grown in culture, MSCs can self-renew, differentiate into several tissues, and modulate the immune response in their surroundings (Dominici et al., 2006; Kuroda & Dezawa, 2014; Uccelli et al., 2008). When activated, MSCs secrete biologically active compounds and generate exosomes that modify the function of their cellular microenvironment (Kou et al., 2022; Madrigal et al., 2014; Phinney & Pittenger, 2017). Due to these characteristics, MSCs are commonly suggested as candidates for cell-based therapy or as the cell source for tissue engineering and cultured meat products. However, it is still unclear how the properties and key biological functions of MSCs *in vitro* are influenced by parameters such as cell source, culture conditions, and environmental factors, including heat stress. Indeed, the long-term epigenetic effects of variations in culture conditions are only beginning to be uncovered (Isik et al., 2021; Sharma & Bhonde, 2020). An example of such an effect is hypoxia-preconditioning, which was shown to significantly reduce global 5hmC in swine MSCs but had no effects on H3K4me3, H3K9me3, or H3K27me3 (Isik et al., 2021). Overall, the long-term effects of stress on MSCs are barely characterized.

Here, we set out to evaluate the immediate and long-term effects of HS treatment on the bovine umbilical cord (bUC)-derived MSC transcriptome. In our previous study, we demonstrated that HS treatment changed the proliferation, differentiation, and immunomodulatory potential of bUC-MSCs (Shimoni et al., 2020). Several HS protocols were shown to have the common effect of reducing proliferation and inducing oxidative stress and premature senescence. However, each HS protocols showed profound variation in gene expression and immunomodulatory potential. Surprisingly, even one hour of treatment at 42°C had a long-term effect on bUC-MSCs function and transcriptional pattern 3 days after the HS, which partially persisted even after eleven passages (about 40 days) in culture (Shimoni et al., 2020). However, at temperatures higher than 41°C the survival rate of mammalian cells decreases with increasing exposure time (Asseng et al., 2021; Moise et al., 2018). Here, to gain an unbiased view of the cell-intrinsic response to heat shock, we performed gene expression profiling using RNA sequencing (RNA-Seq) following exposure of the bUC-MSCs to moderate HS (40.5°C) of different lengths. We hypothesize that transcriptional response to heat stress has two manifestations: a major immediate and transitory reaction of stress response-related genes, and a minor but permanent response of epigenetically altered transcripts. By examining the transcriptional response and functionality of MSCs after mild HS, this work will advance our understanding of the short- and long-term impacts of thermal stress on the organismal stem cell pool.

## Results

### RNA-seq analysis of heat shock treated and control cells

To examine the effects of thermal stress on the cell’s transcriptome, we evaluated the transcriptional response of MSCs to *in vitro* heat shock (HS). MSCs were extracted from the UC of a preterm fetus as previously described (Shimoni et al., 2020). Prior to the HS treatment, MSCs marker expression (Supplementary Figure S1A), proliferation (Supplementary Figure S1B), and differentiation capacities (Supplementary Figure S1C) were examined and validated as in (Nir & Ribarski-Chorev et al., 2022; Shimoni et al., 2020).

HS treatment protocols were designed to examine the short- or long-term effects of exposure to mild HS conditions (Figure 1A). One day after seeding MSCs from early passages (P2–P4), were moved from 37°C to 40.5°C for six-hour (ShortHS) or three days (LongHS) or a six hours HS followed by three days recovery back at 37°C (ShortHS_Recovery). Each HS treatment had a matching 37°C normothermic (NT) control (ShortNT and LongNT). The two time points were selected to capture the immediate transcriptional effects of HS vs. the stable and delayed ones. Immediately after the completion of the six hours or three days treatments, the cells were harvested, viability was checked and pellets were stored at -80° for later matched RNA extraction and sequencing. The viability of the cells and their MSC marker expression levels remained high following the various treatment protocols (Supplementary Figure S1D, S1E). Two or three biological repeats were done for each treatment (Supplementary Table S2). All RNA samples were run together on Nextseq Illumina and showed a high percentage and quality of mapping to exons and low levels of rRNA and mitochondrial DNA contaminations (Supplementary Figure S2A, S2B). Principal component analysis shows that the HS treatment determined the transcriptome profile as samples from the same treatment clustered together and apart from the other treatment groups (Figure 1B). Hierarchical clustering uncovered a secondary factor affecting the transcriptome which could be attributed to the time passed from seeding (Supplementary Figure S2C) or the population doubling (Supplementary Table S3). Several genes were also evaluated by RT-qPCR to validate the sequencing results (Supplementary Figure S2D).

**Figure 1:**
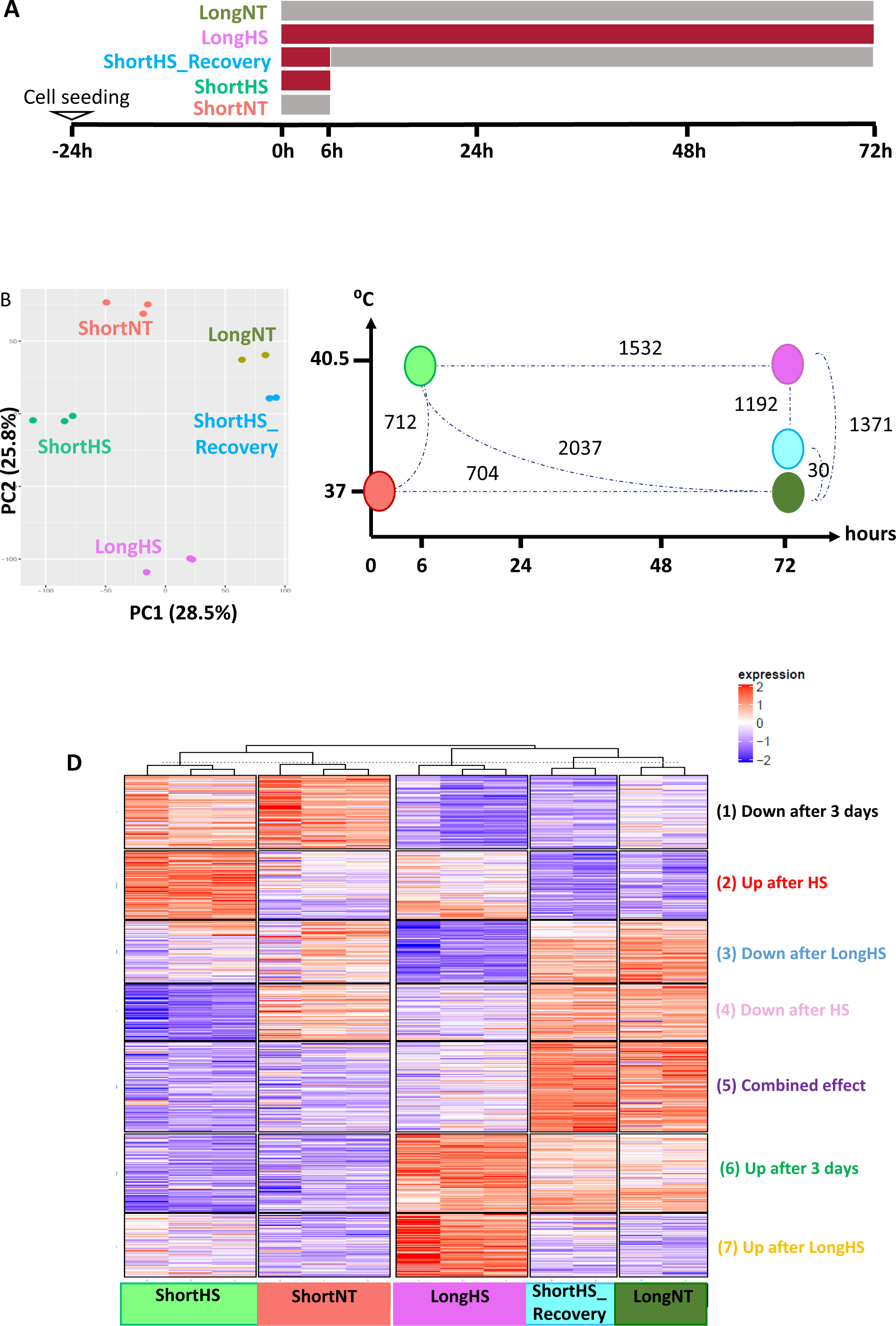
Transcriptome changes in response to heat shock. **(A)** A schematic timeline showing the HS treatments from which RNA-Seq libraries were prepared. MSCs were plated and 24h later exposed to heat shock (HS, 40.5°C) for: 6h (ShortHS), 6h + 3 days recovery at 37°C (ShortHS_Recovery), 72h (LongHS), followed by collection for RNA extraction. Control cells were parallelly plated and maintained in normal temperature (NT, 37°C). Each treatment has colour code which will be used throughout the article: **ShortNT (red), ShortHS (green), LongNT (olive), ShortHS_Recovery (blue), LongHS (purple)**. **(B)** Scatter plot for principal component analysis of global gene expression (RNA-seq) demonstrates clear separation of gene transcripts between groups. **(C)** Number of differentially expressed genes (DEGs) with padj ≤0.05 and log2(FC)≥1 that change between the treatment groups. **(D)** K-means clustering of 3667 DEGs.

We found that the various HS treatments significantly changed the expression levels of a total of 3667 genes (padj ≤0.05 and log2(FC)≥1) (Figure 1C). After six hours of HS, 712 differentially expressed genes (DEGs) were found, whereas after three days of HS 1,371 DEGs were found, most of them downregulated (Supplementary Figure S2E). In the cells that were allowed to recover at 37°C for three days after the short HS, 2,037 genes were changed versus the short HS. However, only a slight difference is found when compared to the long NT sample. This suggests that after a short HS there is a major recovery from the stress, at least on the transcriptional level. The results indicate that short thermal shock affects MSCs’ gene expression rapidly and widely but mostly transiently.

To discern the dominant systematic changes following the different HS treatments, we performed k-means clustering on all 3,667 DEGs (Figure 1D and Supplementary Figure S3). The resulting clusters revealed three major modes of change. Clusters 2 and 4 show the common effect of both long and short HS on the transcriptional pattern. Conversely, in clusters 3 and 7 the change is only observed after the long HS, portraying that more genes were induced after three days, possibly due to a slower transcriptional response rate (Figure 1C). Clusters 1 and 6 show the effect of culturing time on the transcriptional pattern.

While changes in expression patterns are evident from the k-means clustering analysis, we wanted to systematically examine the transcriptional changes following each treatment alone and identify specific biological features and processes.

### Cell cycle and immune response are downregulated as developmental pathways are activated following a long heat shock

To get a comprehensive understanding of all the changes that occurred following the HS, we delved into the analysis of each treatment using the gene set enrichment analysis (GSEA) and gene ontology (GO) tools (as described in the supplementary materials and methods). First, we focused on the changes observed in cells exposed to long HS vs. long NT control (Figure 2A). On the morphological level, cells appeared flattened, with expanded cytoplasm and detectable stress granules (Figure 2B, stress granules marked with arrows). On the transcriptional level, 1,371 DEGs (padj ≤0.05 & log2(FC)≥1) were found (Figure 1C). GSEA analysis show downregulation of cell proliferation and metabolism; namely cell cycle and DNA repair, glycolysis and oxidative phosphorylation (Figure 2C and Supplementary Figure 4A) and upregulated response to stress (Figure 2C). Interestingly, many differentiation and cell-fate related pathways (Figure 2D, Supplementary Figure 4B, C) were also upregulated, as previously suggested (Khan et al., 2020). For example, general terms of cell growth and morphogenesis were enriched, as well as more specific terms like connective tissue, chondrocytes, osteoblasts, kidney, and neuronal system are upregulated. In addition, there is apparent upregulation of cell adhesion molecules, which represents cell-cell interactions and the interaction between the cell and the extracellular matrix (ECM). These molecules promote a broad spectrum of cell signaling that directly or indirectly modulates stem cell proliferation, self-renewal property, adhesion, and multilineage differentiation (Abdal Dayem et al., 2018; Stopp et al., 2013). There is downregulation of gene sets related to inflammation and the immune system, as well as DNA damage and ROS response. This could be explained by the long duration of the HS, which might have required the cells to attenuate their stress response. What is more, these changes were accompanied by the upregulation of post-translational modifiers, specifically histone demethylases and Polycomb group (PcG) complex members (Figure 2E). Additionally, KEGG analysis showed enrichment and upregulation of pathways such as MAPK, PI3K-Akt, RAS and RAP1 (Figure 2F). In general, those pathways are known to regulate cell size, survival, differentiation, adaptation to growth conditions, and stress responses (Li et al., 2017). PI3K-Akt activates the mTOR signaling pathway (Li et al., 2017; Su & Dai, 2017), which is also elevated following heat stress in bovine granulosa cells (Khan et al., 2020). Activation of PI3K-Akt and mTOR pathways plays a central role in cellular senescence and organismal aging and thus acts as a driver of stem cell depletion and reduced tissue regenerative capacity (Antonioli et al., 2019; Johnson et al., 2013; Saxton & Sabatini, 2017).

**Figure 2:**
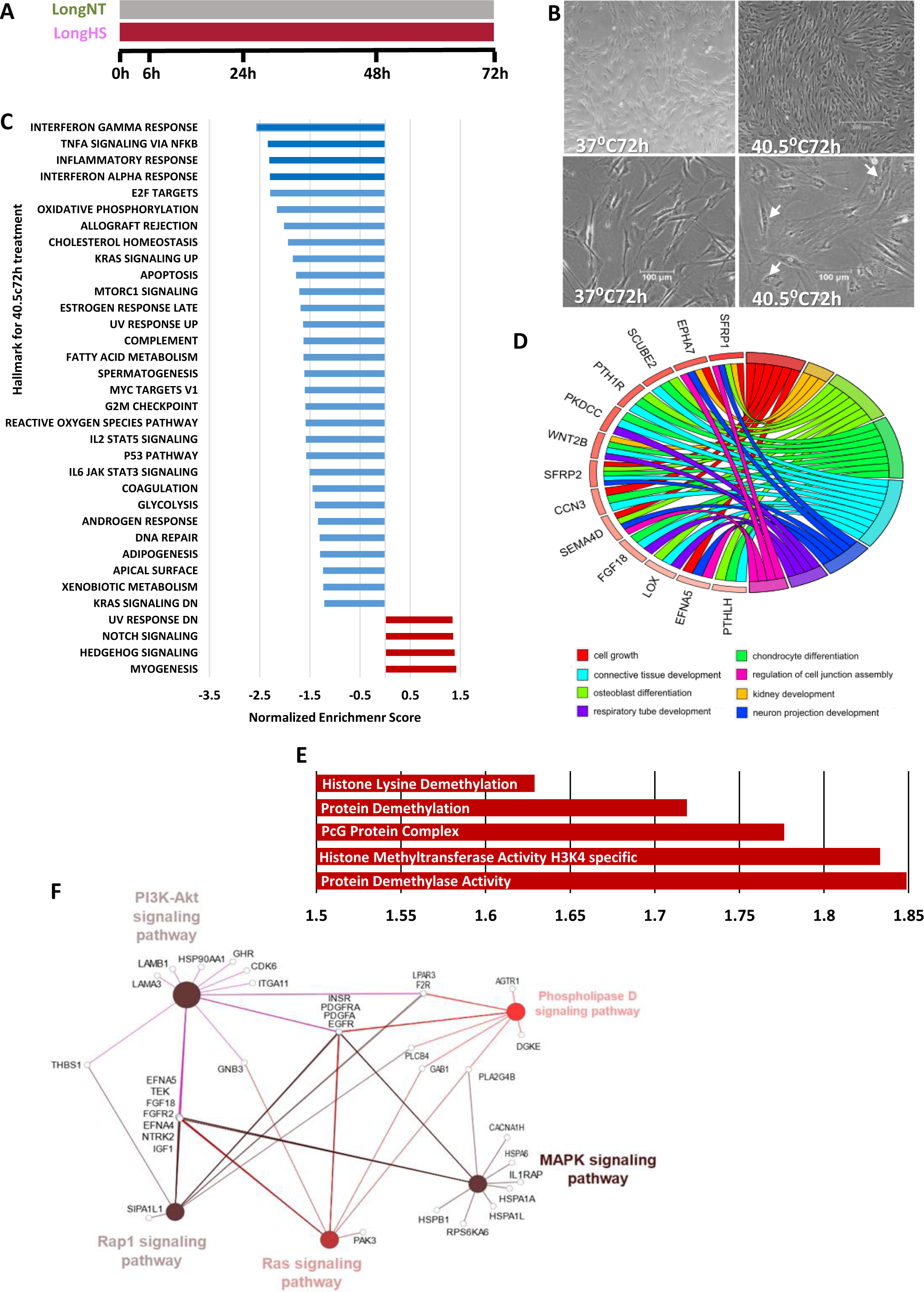
Downregulation of cell cycle and immune response genes while developmental pathways genes are induced following long heat shock. **(A)** Illustration of long heat shock treatment. **(B)** Cells after LongHS get enlarged and flattened. Bright field x4 (up) and x10 (down). Arrows point to stress granules. **(C)** Results of GSEA Hallmark analysis showing enriched gene sets (FDR q-val < 25% and p-val < 0.05). A positive normalized enrichment score (NES) values, mark in red, indicates upregulation in the LongHS phenotype. **(D)** Differentiation pathways significantly upregulated in LongHS vs. LongNT as per g:Profiler analysis. Plot was generated using GOplot R package. **(E)** Upregulated histone modifications and PcG complex following LongHS as per GO analysis with GSEA. **(F)** KEGGs significantly upregulated in differentially expressed genes of LongHS vs LongNT using ClueGo. Colors represented p-value: light red 0.05, red 0.01, brown 0.001. The node size represents the term enrichment significance.

In summary, our data demonstrate that three days of HS treatment are not detrimental to cell viability but have a major effect on cell fate and aging. Although the impact of long HS on differentiation might prove useful as a pretreatment protocol for MSC transplantations, this duration of HS is scarcely experienced in physiological settings. Hence, we wished to examine the effect of short (six hours) HS on the MSCs.

### Stress responses are induced while cellular functions decline following a short heat shock

Initially, we focused on the immediate effects of short HS (Figure 3A) using GSEA analysis. Five out of the ten most upregulated genes in this treatment (HSPA1A, HSPA6, HSPA4L, HSPB1, HSPH1, Figure 3B) were heat stress genes, along with genes that regulate redox maintenance (SLC27A5), insulin levels (MAFA) and mitotic progression (MISP). The most downregulated genes following short HS are involved in RNA interference (RNAi) process (TSNAX), post-splicing multiprotein complex (UPF3A), cell polarity (DAAM2), metabolism regulation (DHRSI2) and chromatin and transcription process (H1-2). These genes are mostly included in the Hallmark gene sets that are enriched for up- and down-regulated processes following six hours of HS (Figure 3C and Supplementary Figure 5A, 5B). Short heat stress induced stress response and differentiation pathways while halting oxidative phosphorylation and cell cycle. Upregulated biological processes enriched in GSEA were stress response, autophagy, development, and mRNA transcription (Figure 3D, Supplementary Table S4), while downregulated processes were all related to DNA replication (Figure 3E and Supplementary Figure 5B). In addition to GSEA, we performed the analysis in g:Profiler, KEGG and Reactome which also suggests that stress response and ER activity were induced whereas DNA replication, with an emphasis on initiation, was decreased (Figure 3F). After discovering dramatic short-term changes, we proceeded to check how many of these changes remained stable. To elucidate these stable changes, the treated cells were allowed to recover in normothermia for three days before the transcriptional analysis.

**Figure 3:**
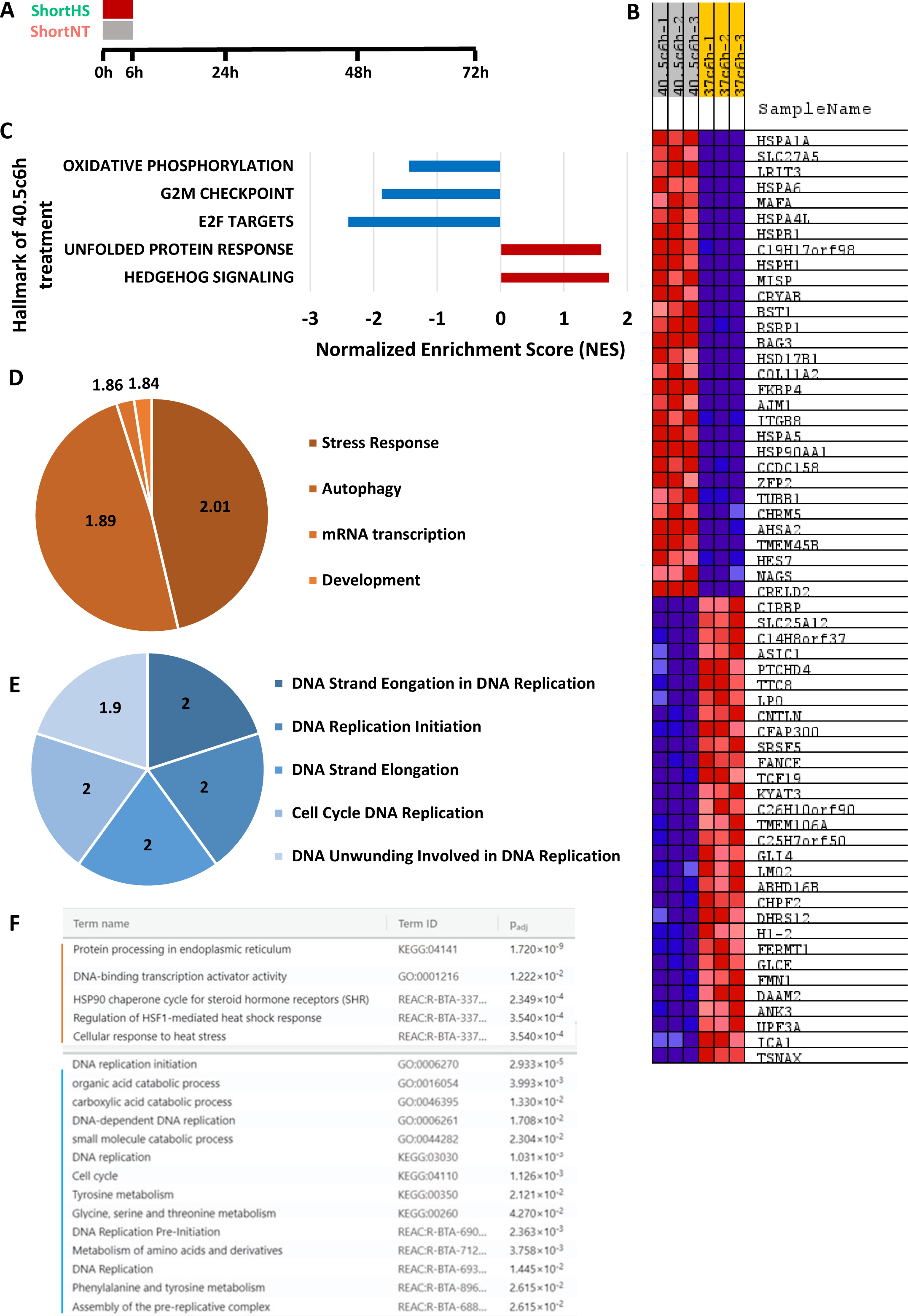
Induced stress response and reduced cellular functions following short heat shock. **(A)** Illustration of short heat shock treatment. **(B-E)** Gene set enrichment analysis (GSEA) results of 40.5c6h vs. control comparison. RNA-Seq was performed on samples collected after 6 hours at 40.5c compared to cells cultured in 37c (control). **(B)** Heat map of the top 30 marker genes upregulated and downregulated in short heat stress. Expression values are represented as colors and range from red (high expression), pink (moderate), light blue (low) to dark blue (lowest expression). **(C)** Results of GSEA Hallmark analysis showing enriched gene sets (FDR q-val < 25% and p-val < 0.05). A positive normalized enrichment score (NES) values, marked in red, indicate enrichment in the 40.5c6h phenotype. **(D)** Upregulated GSEA biological processes significantly enriched (FDR q-val < 0.25 and p-val< 0.01) in the comparison of ShortHS vs. ShortNT, ordered by the increasing NES (dark color is the most enriched). Due to high number of GOs, the GOs with similar process were combined under one general phrase, which is represented by one part of the pie chart (Supplementary Table S4). The number defines average normalized enrichment score. **(E)** Downregulated GSEA biological processes significantly decreased (FDR q-val < 0.25, p-val < 0.01) in the comparison of ShortHS vs. ShortNT, ordered by the increasing NES. **(F)** Functional enrichment analysis of the upregulated (red) and downregulated (blue) differentially expressed genes (DEGs) between ShortHS and ShortNT cells using g:Profiler.

**Figure 5:**
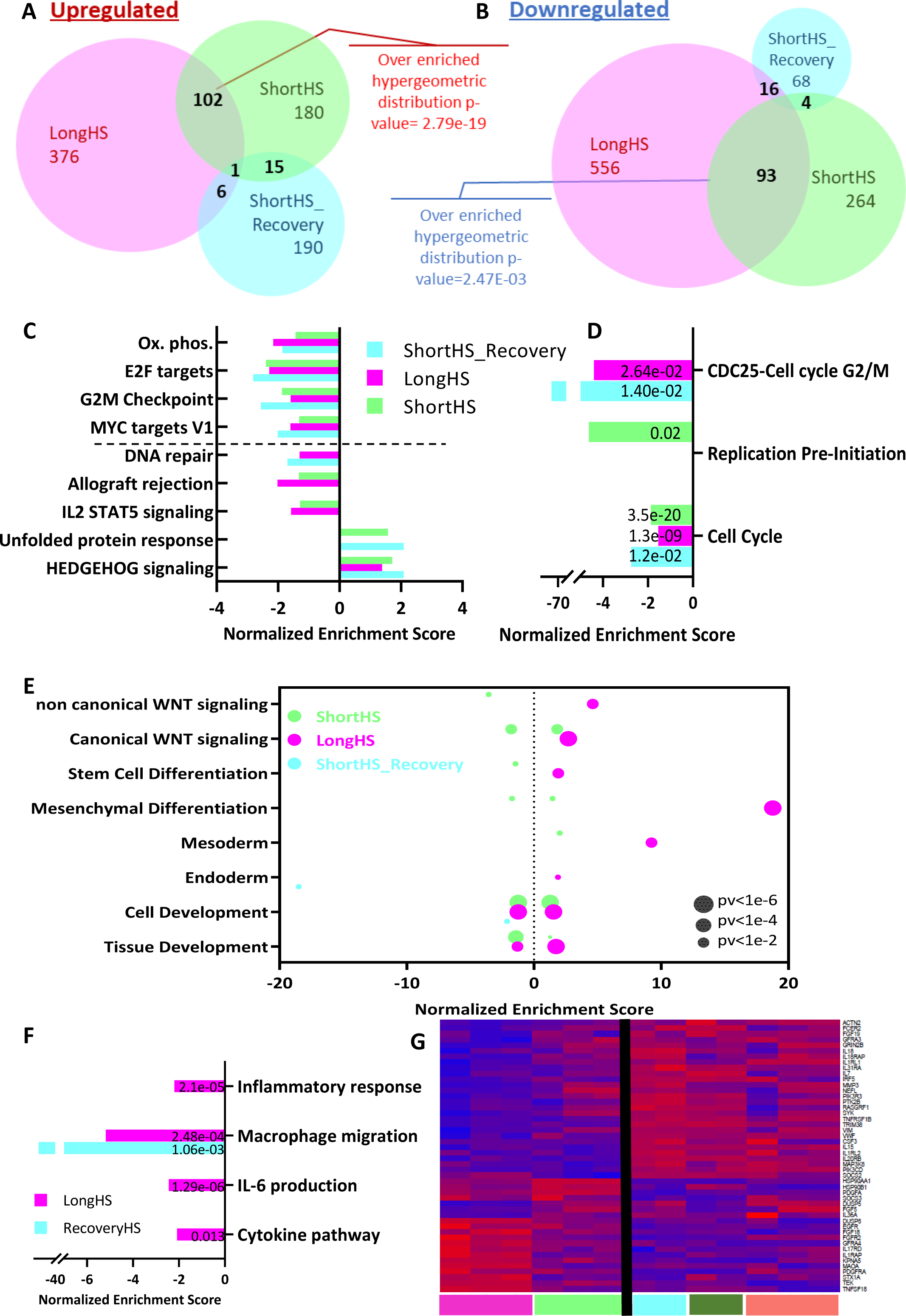
Cell cycle and metabolism are commonly reduced following all heat shock protocols, while other processes related to key characteristics of MSC are differently regulated. **(A)** Venn diagram of upregulated genes (padj < 0.05) between the treatment groups. **(B)** Venn diagram of downregulated genes (padj < 0.05) between the treatment groups. **(C)** Heat shock common hallmarks and their enrichment score. Above hyphenated line are hallmarks common to all 3 HS groups. **(D)** Cell cycle related biological processes, effected by HS, and their enrichment score. N00455: CDC25-Cell cycle G2/M (KEGG pathway), GO:0007049: Cell cycle (MGI database), R-BTA-69002 – DNA Replication Pre-Initiation (Reactome). **(E)** Enrichment of biological processed related to development effected by HS. GO:0060070: Canonical WNT signalling, GO 0048863: Stem cell differentiation, GO:0048468: Cell development, GO:0009888: Tissue development, GO:0007492: Endoderm development. GO:0007498: Mesoderm development, GO:0048762: mesenchymal cell differentiation. **(F)** Enrichment of biological processed related to immune system effected by HS. GO:0002526: Acute Inflammatory response, GO:1905517: Macrophage migration, GO:0032635: IL-6 production, Cytokine pathway list was taken from PathCards v5.7.551.0. **(G)** Cytokine pathways genes that are differentially expressed in LongHS vs. LongNT.

### Long term effects of short heat stress

To identify the transcriptional changes which occurred after short HS but remained stable even after 3 days of recovery, we compared the three different time points: control (as the zero-time point), six hour HS and six hour HS followed by three days of recovery (ShortNT vs. ShortHS vs. ShortHS_Recovery, Figure 4A). Thus, we re-analyzed the data (see Supplementary materials and methods) using K-means clustering to visualize transcriptomic differences (Figure 4B). While after recovery most genes regained similar expression levels to that of the control sample, two clusters in which the differential expression persisted were identified. Cluster 2 genes were lowly expressed in the control sample but were then upregulated after the short HS and remained active after recovery. GO annotations and KEGG pathways enriched in this cluster are mostly related to stress and protein degradation (Figure 4C). This implies that some elements of the stress response to acute stress last for more than three days. Cluster 3 is the mirror image of cluster 2, presenting stably downregulated genes related to proliferation and metabolism but also senescence and DNA damage (Figure 4D). This might suggest that following short HS, the cells enter a state of quiescence to maintain their stemness. This data is in agreement with our population doubling time analysis, which shows slower proliferation after both the long and the short HS (Figure 4E).

**Figure 4:**
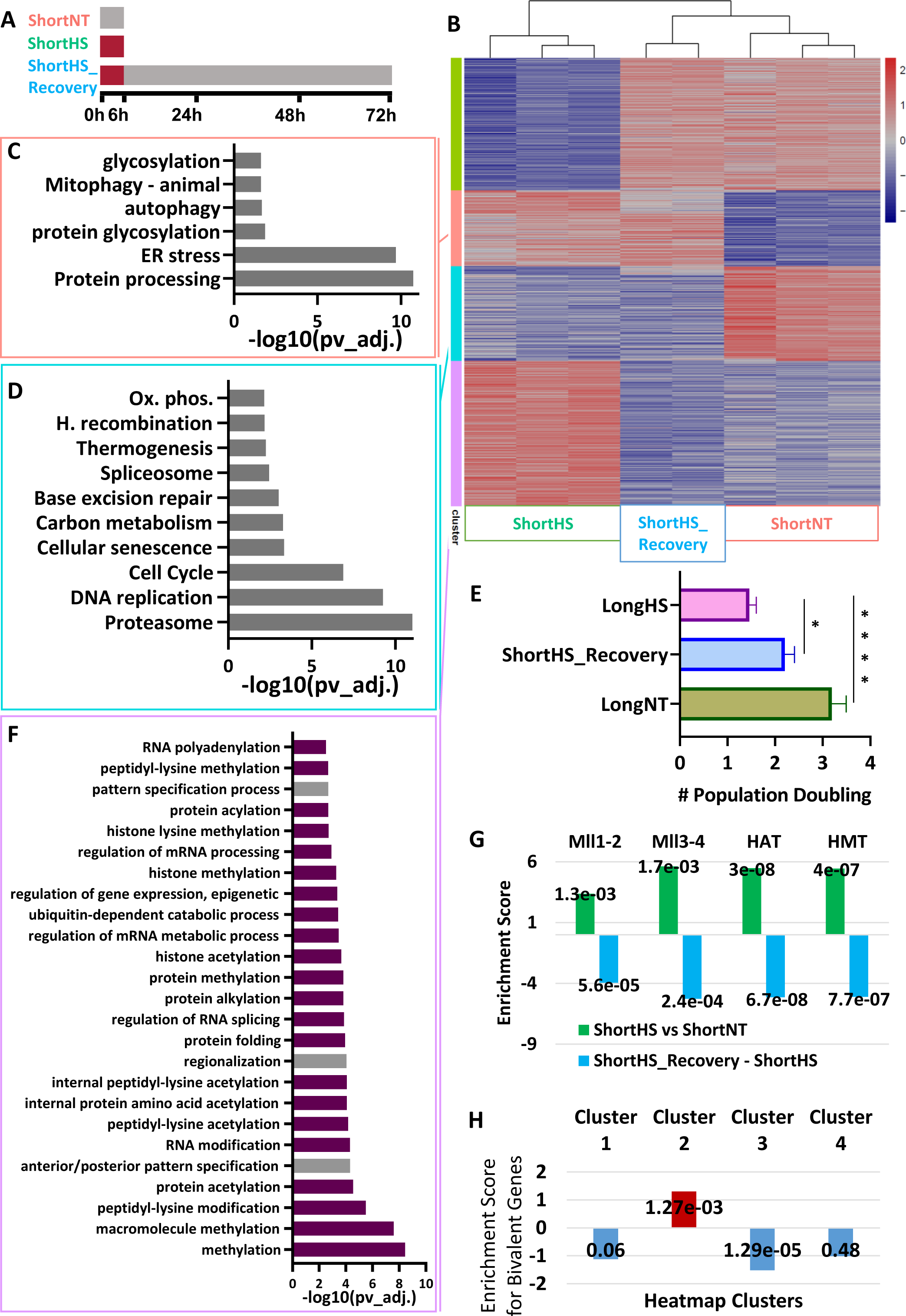
Transient upregulation of chromatin modifiers after short heat shock and stable activation of lineage commitment genes, maintained even after three days recovery. **(A)** Illustration of ShortHS_Recovery vs ShortHS treatment. **(B)** Heat map of genes differentially expressed (padj < 0.05) between ShortHS and ShortNT. **(C)** Biological processes enriched in cluster 2 (pink) **(D)** Biological processes enriched in cluster 3 (blue) **(E)** Population doubling time for prolonged treated groups. **(F)** Biological processes enriched in cluster 4 (purple) **(G)** MLL (GO:0044665, GO:0044666), HAT (GO:0000123) and HMT (GO:0035097) are upregulated following ShortHS but return to normal/downregulated during recovery period **(H)** Bivalent genes are upregulated following ShortHS and remain in this state even after recovery period.

Similar analysis done on the other two clusters, 1 and 4, raised an interesting point. Of the most 25 significantly enriched GO terms in cluster 4, only 3 are not related to transcriptional regulation, mostly epigenetic modification (Figure 4F). This raised the intriguing hypothesis that the epigenetic landscape is altered following the short HS, possibly with long term consequences. To examine how a short period of stress could have that kind of persistent effect, we used the EpiFactors Database (Medvedeva et al., 2015) and related MGI annotations to identify enriched epigenetic modifiers in the data. Several epigenetic complexes were found highly enriched in Cluster 4, i.e., upregulated after six hour HS and then back to low expression after recovery (Figure 4G). Interestingly, those complexes are all related to the mixed lineage leukemia (MLL) proteins which catalyze the trimethylation of histone H3 Lys 4 (H3K4me3), a mark associated with active or poised transcription and found in the promoters of most active or poised genes (Harikumar & Meshorer, 2015). Bivalent genes are lineage-specification genes that carry both H3K4me3 and H3K27me3 histone marks and are regulated by the balance between the two histone marks. This balance is mediated by PcG members and the MLL coactivator complex (Noer et al., 2009). We searched for known targets of the MLL complex in our list and found cluster 2 (i.e. upregulated in both short- and long-term) to be significantly enriched for bivalent genes (Mas et al., 2018) crucial to cell fate regulation: several FGF family members, FZD3, JUN, FOXF1, VEGFA and KDM6B, to state a few (Supplementary Figure S6). This was in contrast to the other clusters, where these and other bivalent genes were significantly underrepresented (Figure 4H). Overall, this data sheds light on the long-term effects of short HS and suggests that a short stress event can modulate the epigenetic regulation of key cell fate genes.

Next, we wished to uncover the transcriptional changes shared by all HS-treated cells.

### Key characteristics of MSC vary with heat stress duration

So far, we have shown considerable effects of short and long HS *in vitro.* To examine to what extent the DEGs are shared between the timepoints, we had to eliminate the effect of culturing time and the circadian clock and focus solely on the effect of HS. To that end, we removed all DEGs that changed between the short and long NT controls (see Supplementary materials and methods for the analysis) and examined which DEGs were commonly upregulated (Figure 5A) or downregulated (Figure 5B) in all HS. A total of 102 upregulated and 93 downregulated DEGs were shared by short and long HS treatments, a highly significant enrichment over what could be expected by chance. While the commonly upregulated genes were mostly related to stress response, the commonly downregulated genes were not significantly enriched to any specific pathway or process. Hence, we examined if the different DEGs are annotated to shared biological processes and pathways which are therefore affected by both short and long HS. Important cellular functions like oxidative phosphorylation and cell cycle were found to be impaired in all HS; immune system was downregulated, and the hedgehog signaling pathway was significantly upregulated following HS, even after the recovery period (Figure 5C, Supplementary Figures S7A-D).

To understand the effect of HS on the biological processes related to the key capacities of MSCs, namely self-renewal, differentiation and immunomodulation, we compared the relevant biological processes and functions between different treatments. For cell cycle we saw that while in all three treatments the downregulated DEGs were enriched for the general term ‘cell cycle’, only the short HS is downregulating specifically DNA replication pre-initiation (e.g., genes like MCM2 and 7, CDC6 and E2F1). The three days samples showed downregulation of genes required for the G2/M phase (namely cyclin A and B and CDC25), suggesting a block before mitosis (Figure 5D). Interestingly, the cells that got the short HS but were let to recover for three days in normothermia show both the G1/S and the G2/M effects although to a lesser degree, suggesting that even short thermal stress might disrupt cell cycle regulation and proliferation. These results are supported by the population doubling we saw in the culture following the experiments (Figure 4E) as well as in our previously described cell cycle analysis (Shimoni et al., 2020).

As for differentiation, it appears that several developmental pathways were activated, depending on the heat shock treatment (Figure 5E, Supplementary Figure S7E-F). The long HS induced various differentiation pathways; the most enriched were the differentiation to mesoderm/mesenchymal fate. Notably, both canonical and non-canonical WNT pathways were upregulated after long HS, suggesting a general shift in cell identity. The short HS upregulates the canonical WNT but downregulates the non-canonical (WNT5a related) pathway (Figure 5E, Supplementary Figures S7E, F). In line with the fact that KEGG definitions like “canonical WNT pathway” includes both activators and suppressors, this and other pathways are enriched in both the down and upregulated DEGs. Interestingly, HOX genes related to skeletal morphogenesis and several epigenetic factors (e.g., Setd2, Kdm6a, Jarid1) were upregulated following both long and short HS; suggesting that heat shock treatment may slow proliferation while promoting MSC’s commitment and differentiation.

Immunomodulatory properties of MSCs have previously been shown to decline following stress (Bagath et al., 2019; Shimoni et al., 2020). Major downregulation of immune related pathways was detected after HS (Figure 5F). GO analysis in GSEA revealed considerable effects on the production of interleukins (ILs), TNFs and INF-ɤ. In addition, we observed the downregulation of antigen processing, acute inflammatory response, response to chemokines, and the migration and chemotaxis of macrophages, neutrophils, and granulocytes (Supplementary Table S5). The list of 760 genes in the "Cytokine Signaling in Immune system SuperPath" (https://pathcards.genecards.org) was analyzed and 38 of them (for example IL1Rs, IL7 and IL18) were downregulated after long HS while 17 (like FGF2 and FGFR2 which are related to differentiation (Khalid et al., 2021; Zhang et al., 2022)), are upregulated (Figure 5G). Overall, genes related to IL production are significantly downregulated (hypergeometric p-value = 2.49e-04, Figure 5G).

## Discussion

Here, we show that the transcriptional changes following heat shock *in vitro* are broad and can alter every functional aspect of MSC identity. We analyzed RNA-seq from bUC-MSCs in normothermia vs. short or long exposures to HS, to investigate the subsequent changes in the transcriptional landscape that might lead to a shift in the cell’s identity and fate. We were particularly interested in identifying the long-term effect of thermal stress on cell proliferation, differentiation, and immunomodulation capacities; as those key MSC features were amongst the most abundant GO term groups. To examine the relevance of our data to physiological heat stress, we compared our DEGs to a list of 55 genes defined as thermal stress responsive in a bovine study done *in vivo* (Fang et al., 2021): 36 of the 55 genes were found to be differentially expressed following short, long or both HS groups. The genes EIF2A, HSPA1A, HSP90AA1, and HSF1 were considered by the authors key genes that responded to thermal stress of Holstein dairy cattle. From our experiments, HSPA1A, and HSP90AA1 were indeed upregulated DEGs for short and long HS groups, and EIF2A for shortHS group only. HSF1 was not differentially expressed in our treatments, maybe due to our use of 40.5°C which is mild relative to the 42°C used by the authors. In addition, Fang et al. bioinformatics analyses pinpointed biological processes and pathways associated with thermal stress that are very similar to those we have identified, e.g., protein folding, transcription factor binding, immune effector process, negative regulation of cell proliferation, PI3K-Akt signaling pathway, and MAPK signaling pathway.

The immediate transcriptional changes found following short HS include the onset of several cellular stress responses and cell cycle arrest in G1/S. This checkpoint arrest is a known adaptive cellular response to heat stress, in which cells slowdown proliferation rate while rapidly upregulate the transcription and translation of HSPs to maintain cell homeostasis and retain their cellular functions (Richter et al., 2010). Our previous results, which showed cell cycle arrest at G1/S after short HS (Shimoni et al., 2020) also support this finding. Two possible interpretations arise, each with a different implication for stemness and differentiation: in one, following the short HS the cells enter a quiescent state; this is in alignment with the observed metabolic changes (reduced amino acid metabolism, reduced oxidative phosphorylation, increased glycolysis, and mitophagy induction of mitochondrial renewal, etc.), as reviewed in (van Velthoven & Rando, 2019). This suggest that short HS treatment encourages the retention of MSC stemness. In the other interpretation, a long G1 may allow for the accumulation of the epigenetic changes needed for the initiation of fate decisions (Dalton & Coverdell, 2015). This interpretation might explain the transient upregulation of many epigenetic factors and the stable changes in many lineage commitment genes observed after three days recovery. To mechanistically challenge those two interpretations and to determine whether HS promotes stemness or differentiation, further study, preferably on the single cell level, is required to mechanistically challenge these two interpretations.

Evident changes were observed in the morphology and proliferation rate of the cells three days after the Short or Long HS treatment. However, on the transcriptional level, the majority of ShortHS DEGs reverted to normal expression levels after three days recovery, and only a subset of the DEGs remained. In this subset of stable DEGs we see the upregulation of several developmental genes known as "bivalent" for their dual histone lysine tri-methylation marking in both K4 (active mark) and K27 (repressive mark). Those genes are known targets of the MLL complex, which carries out the methylation of histone H3 lysine 4 and was transiently overexpressed after the initial HS event. So, a sequence of events can be hypothesized. First, the short HS upregulates the MLL complex members, thus shifting the balance between the active and repressive histone marks. Consequently, the chromatin of the previously poised (and silent) bivalent genes opens to enable transcription of developmental genes, priming the cells toward cell fate determination. Evaluation of this hypothesis will require follow-up studies examining changes in epigenetic marking following HS.

The idea that cells after HS pretreatment are more prone to differentiation is tempting. If indeed priming MSCs by specific stress can help us direct their fate, it could provide us with another tool in the cellular therapy toolbox. Moreover, as the idea of cellular agriculture, or cultured meat, gains increasing attention, more cost effective and clean ways to differentiate the source cells toward the intended fate (usually muscle or fat) are needed. Current differentiation protocols require large amounts of expensive and unstable growth factors, which raise costs and hinder commercialization of cultured meat. Every pretreatment protocol that is easy, cheap, and reduces the time and resources necessary for differentiation, will have huge economic and environmental consequences.

On the clinical level it is interesting to note the negative effect of long HS on MSC immune activity, possibly compromising the cells’ immunomodulatory functions. Indeed, reduced immunomodulation was previously observed when we co-cultured MSCs subjected to HS with macrophages (Shimoni et al., 2020). The reduced immunomodulation capacity of the stressed MSCs might account for the increased production of inflammatory cytokines found in mammary epithelial cells following HS (Gao et al., 2019). Hence, the increased rates of inflammatory diseases in dairy cows during the summer months could be partially attributed to the failure of the malfunctioning MSCs to modulate the inflammatory response of the surrounding cells to HS. Overall, identifying the long-term effect of HS on MSC capacities suggest an explanation for the seeming contradiction between the in vitro experiments which show beneficial effects to HS priming and the in vivo data that demonstrate harmful physiological consequences to thermal stress (reviewed by (Roth, 2020)).

Although we have demonstrated that HS affects MSCs’ transcriptome, our study has some limitations. Population heterogeneity, circadian clock, culturing time, and culture conditions all have effects on RNA-seq results that need to be controlled for (Supplementary Figure S8). To counter this drawback we (1) made sure that at the end of the experiment (either six hours or 3 days) all the treatment groups had similar culture density (Supplementary Table S2) (2) sequenced two or three independent biological repeats from each treatment group (Supplementary Table S3) (3) to reduce heterogeneity that might originate in genetic background, the same MSC line was used in all three repeats. To counter the effects of culturing time, we used untreated (NT) controls, one for the short HS, and the other for the long HS. When we compared gene expression at the short and long time points, all DEGs that were apparent between the NT control time points (and were assumed to be the result of culturing effect) were ignored. Interestingly, two transcription factors regulating the circadian clock, BHLHE40 and TIMELESS, were found among those ignored DEGs. These two transcription factors together regulate the expression of 202 other DEGs in this group (Supplementary Figure S8B). These oscillating genes may partly explain the relatively high number of DEGs due to culturing effect since the ShortHS and ShortNT samples were taken at noon (24h after seeding + six hours treatment) while the three day samples (LongHS, LongNT and ShortHS_Recovery) were collected in the morning (exactly 96h after seeding).

## Conclusions

The use of MSCs for cell therapy or cultured meat has several advantages, such as high availability, low production price, and fast and easy differentiation. Nonetheless, our limited understanding of their heterogeneity, fate determination, and immunomodulation holds back further MSC applications. It is therefore essential to improve the consistency and efficacy of MSCs. Such enhancement can be achieved using in-vitro preconditioning treatments such as HS (Choudhery, 2021). Here we show that the correct application of HS allows for the enhancement of desired MSC characteristics and induction of a wide range of stem cell fates and differentiation pathways.

## Materials and Methods

### Cell Culture

Bovine mesenchymal stem cells were isolated, cultured and characterized based on generally accepted criteria (Dominici et al., 2006; Uccelli & al., 2008) as we have previously reported (Nir & Ribarski-Chorev et al., 2022; Shimoni et al., 2020). Briefly, umbilical cords of Holstein dairy cows were obtained from abattoirs located in the north of Israel. In the lab, the umbilical cords were digested and plated as previously described in (Toupadakis et al., 2010), followed by expansion in low glucose Dulbecco’s Modified Eagle’s Medium (Gibco, 31885-023 or Biological Industries (BI), 04-001-1A), containing 10% Fetal Bovine Serum (FBS) (BI, 04-001-1A), penicillin-streptomycin solution 1% (BI, 03-031-1B), Glutamine 1% (BI, #03-020-1B) and cryopreserved at different passages using FBS with 10% DMSO (Sigma-Aldrich, W387520). The media was changed every 2-3 days and all cells were cultured in a humidified incubator with a controlled environment of 5% CO2 and a temperature of 37°C, unless mentioned otherwise.

### Cell Characterization

Three bovine MSCs lines were used in this study. Two were previously examined and verified (Nir & Ribarski-Chorev et al., 2022; Shimoni et al., 2020) and the third was examined and verified in this study (Supplementary Figure 1 (S1A-S1C)) before subsequent heat shock experiments. One line was used for the RNA-seq experiment, while all three lines were used for RNA-seq validation and further examinations. All the experiments described were done on cells in passages 2-4, and at least 3 biological replicas were used unless specified otherwise.

### Cell Death Quantification Using Propidium Iodide (PI)

For quantification of cell death in culture, cells were harvested by trypsinization, washed with PBS, and re-suspended in 10 g/mL PI for 5 min on ice. The percentage of live/dead cells was determined within 1 hour of staining by flow cytometry (CytoFLEX, Beckman Coulter, Indianapolis, IN, USA). 10,000 events were collected per sample. At least three biological repeats were used for each treatment. Data analysis was performed using Flow-Jo software.

### Heat-shock Induction

MSCs at early passages (P2–P4) were seeded in 6-well plates (Costar, 3516) at different concentrations to allow similar confluence at the end of the experiment, as follows:

| Time of treatment | Temperature | # of cells seeded | abbreviation |
| --- | --- | --- | --- |
| six hours | 37°C | 250,000 | ShortNT |
|  | 40.5°C | 250,000 | ShortHS |
| Three days | 37°C | 42,000 | LongNT |
|  | Six hours at 40.5°C, three days recovery at 37°C | 72,000 | ShortHS_Recovery |
|  | 40.5°C | 72,000 | LongHS |

The cells were incubated for 24 hours in normal conditions before treatment onset (time 0).

### Cell Counting

Following the HS, the cells were washed with Dulbecco’s Phosphate Buffered Saline (BI, 02-021-IA) and trypsinized (BI, 03-053-1A). 6μL of trypsinized cells were mixed with 6μL of trypan-blue (Sigma, T8154), loaded into an appropriate slide, and counted using an automated cell counter (TC20, Bio-Rad Laboratories Hercules, CA, USA). Three to five biological repeats were performed.

### RNA-sequencing

Total RNA from the heat shocked cells was extracted via PureLink RNA Minikit (Invitrogen, 12183018A). RNA concentrations were measured using a NanoDrop ND-1000 spectrophotometer (NanoDrop Technologies) and the quality of extracted RNA was confirmed (RNA integrity number (RIN) > 9) using an Agilent 2100Bioanalyzer before further processing. Library preparation was done using the INCPM mRNA library of the Weizmann Institute, Rehovot. The quality of cDNA libraries was determined using a tape-station. Sequencing was done using Nextseq Illumina in the INCPM of Weizmann Institute, with single-end reads of 75bp. Sequencing was done on 14 samples from the same MSC line as in (Shimoni et al., 2020), and after initial quality analysis one sample was removed and we were left with 13 samples to perform the in-depth analysis: 3 biological replicas for ShortNT, ShortHS and LongHS treatments, and 2 biological replicas for LongNT and ShortHS_Recovery treatments.

### Data Analysis

Following sequencing, reads were trimmed using Cutadapt, mapped to genome (Bos_taurus.ARS-UCD1.2) and counted using STAR and HTSeq. Further quality control of alignment sequencing data was done using the Qualimap. Normalization of the counts and differential expression analysis were performed using DESeq2. Hierarchical and K-means clustering as well as data visualizations were performed using R packages. Gene names were taken from the Ensemble BioMart ARS-UCD1.2 dataset. Gene Ontology (GO), KEGG and Reactome pathways enrichment analyses were conducted using both ranked gene list analyses using Gene Set Enrichment Analysis (GSEA) software version 4.1.0 and DEGs list analysis using g:Profiler version e103_eg50_p15_eadf141 and ClueGO v2.5.7 (Cytoscape plug-in) with default parameters. KEGG pathway maps were downloaded from KEGG pathway database. String datasets were used for functional protein association networks. Epigenetic complexes analysis was performed using the EpiFactors Database (Medvedeva et al., 2015) as well GO annotations of the MGI database (Mouse Genome Informatics). The cytokine pathway list was taken from PathCards v5.7.551.0. Bivalent genes enrichment was done by comparing our list to “3,868 bona-fide bivalent genes” in supplementary table No. 1 of (Mas et al., 2018). For detailed information see Supplementary Materials and Methods.

### RNA extraction, Reverse transcription, and Real-time PCR

RNA extraction from cells was carried out using GenElute™ Total RNA Purification Kit (Sigma, RNB100-50RXN). RNA was then reverse-transcribed into cDNA using qScriptTM cDNA Synthesis Kit (Quanta-bio, 95047-100). Real-Time PCR (RT-qPCR) reactions were performed using Fast SYBR Green Master Mix (Applied Biosystem, 4385614) in an ABI Step-One Plus Real-Time PCR system. All primers were verified by standard curve evaluation and are shown in Supplementary Table S1. Primers for expression analysis were designed on exon-exon junctions and a -RT control was performed each time. Relative mRNA fold change was calculated with the ΔΔCT method, using 1-2 control genes (PSMB2 and RPS9) as reference.

### Statistical Analysis

GraphPad Prism (La Jolla, CA, United States) was used for statistical analysis and visualization of RT-qPCR. p < 0.05 was considered statistically significant. For enrichment analysis hypergeometric p-value was calculated using The Graeber Lab online calculator 2009 © (https://systems.crump.ucla.edu/hypergeometric/). See Supplementary Materials and Methods for more detailed information.

### Availability of data and materials

The datasets supporting the conclusions of this article are available in the GEO repository, GSE214467 study at: https://www.ncbi.nlm.nih.gov/geo/query/acc.cgi?acc=GSE214467

Some datasets supporting the conclusions of this article are included within the article additional files.

## Acknowledgments

We are grateful to all the members of the Schlesinger lab for their ideas and support; Prof. Z. Roth for scientific discussion, encouragement and support; Joseph Kippen for critical reading and reviewing; the Weizmann Institute Next Generation Sequencing Facility, for sequencing support; The authors also thank Dr. S. Shainin and Mr. S. Yaakobi from the Volcani Center for help with obtaining the umbilical cords.

## Authors contributions

IRH: conception and design, designing and performed the experiments, bioinformatics analysis and data interpretation, and manuscript writing. GS: validation of MSC lines by flow and RT-qPCR. CS: support with MSC extraction and culturing. SS: conception and design, assembly of data and data analysis, manuscript writing. The authors read and approved the final manuscript.

## Ethics approval and consent to participate

The procedures of isolated UC-MSCs were conducted abiding by the Institutional Animal Care and Use Committee of the Israel Ministry of Agriculture and Rural Development, permit #11380.

## List of abbreviations

MSCs: Mesenchymal stem cells
bUC-MSCs: Bovine umbilical cord-derived mesenchymal stem cells
ER: endoplasmic reticulum
FBS: Fetal Bovine Serum
RNA-seq: RNA sequencing
RT-qPCR: Reverse transcriptase-quantitative polymerase chain reaction
HS: Heat shock
NT: Normothermic
DEGs: Differentially expressed genes
GSEA: Gene set enrichment analysis
GO: Gene ontology
ECM: Extracellular matrix
PcG: Polycomb group
MLL: Mixed lineage leukemia
H3K4me3: The trimethylation of histone H3 Lys 4
ILs: Interleukins

## Supplementary figure legends

**Supplementary Figure 1:** Characterization of the bUC-MSC line used for RNA-seq validation. (A) Expression of positive (cd29, cd44, cd 90) and negative (cd45) MSC markers normalized to RPS9. (B) Population doubling time, 3-4 biological replicas at each time point. (C) MSCs differentiate to chondrocytes (Alcian blue staining), osteoblasts (Alizarin red staining) and adipocytes (BODIPY™ 493/503 and DAPI staining). Chondrocyte and osteoblast staining was observed using EVOS® FL Auto microscope scale bar 100m. Adipocyte staining was observed using a fluorescent microscope, scale bar 50m. (D) PI staining for live/dead cells after all HS treatments (E) Positive and negative markers of MSC, normalized to RSP9 housekeeping gene, after all HS treatments

**Supplementary Figure 2:** RNA-seq quality. Bio detection histogram (Qualimap application) showing which kind of features are being detected in control **(A)** vs. HS **(B)** groups, similar distribution was seen in all samples. The x-axis shows all the groups included in the annotations file. The gray bars are the percentage of features of each group within the reference genome. The striped color bars are the percentages of features of each group detected in the sample with regards to the reference. The solid color bars are the percentages that each group represents in the total detected features in the sample. **(C)** Hierarchical clustering of the samples presents clustering by treatments and by the time the cells were in the cutler (culturing effect). **(D)** Validation of RNA-seq by RT-qPCR. Raw read counts were used for sequencing data plots. **(E)** The direction of DEGs regulation per treatment, shows more genes are downregulated during heat shock than upregulated.

**Supplementary Figure 3:** Transcriptional changes in differentially expressed genes of MSC following heat shock. **(A)** K-means clustering of 3667 DEGs, with main biological processes in each cluster. **(B-H).** Focusing on DEGs enrichment for each treatment pair. Each graph: In the top left there is a barplot for the number of DEGs in each cluster. The main graph is a scatterplot of the RNA-seq enrichment. Axes colours match treatment colour coding [ShortNT (red), ShortHS (green), LongNT (olive), ShortHS_Recovery (blue), LongHS (purple)]. Each axis represents upregulated DEGs of specific treatment as specified on the axes. DEGs coloring matches the color code of clusters and biological processes in figure A, right side. Non-differential genes are gray.

**Supplementary Figure 4:** LongHS halts cell cycle and promotes cell developmental fate. **(A)** Cell cycle arrest during LongHS (KEGG Pathway bta041100). Upregulated (pink) and down (yellow) DEGs (padj <0.05 and log2(FC) >1) of LongHS treatment are marked. On right side, RT-PCR validation of cell cycle genes, normalized to PSMB. (B) Upregulated DEGs marked in red of adhesion molecules (KEGG map bta04514) following LongHS. (C) Upregulated DEGs marked in red of axon guidance (KEGG map bta04360) following LongHS.

**Supplementary Figure 5:** ShortHS halts DNA replication. **(A)** Genome viewer (IGV) screenshot of genes that become significantly up-regulated following HS treatments. FOXF1-transcription factor associated with cell differentiation; AHSA2 - activator of HSP90. **(B)** DNA replication is downregulated during ShortHS (KEGG map bta03030). Downregulated (blue) DEGs (padj <0.05 and log2(FC) >1) are marked.

**Supplementary Figure 6:** Upregulated bivalent genes following ShortHS that maintain direction of regulation during recovery period. Biological process (BPs) enriched for bivalent genes found in Cluster 2 of Figure 4B, lowly expressed genes in the control sample that were upregulated after the short HS and remained active during recovery. (BP’s term size used in g:Profiler was between 5-1000; Figure prepared using GOChord plot in RStudio). On right is BPs with color coding as per the legend. From left are genes common to at least 4 BPs, arranged by log2 Fold Change.

**Supplementary Figure 7:** Cell functions modified by heat shock. **(A)** Hallmarks affected by specific heat shock treatments. ShortHS has no specific process, only ShortHS_Recovery, and LongHS groups. **(B)** All genes with padj<0.05 and no culturing effect that are affected by HS treatments in the cell cycle (KEGG Pathway bta041100). Blue and red, downregulated and upregulated genes accordingly.

**Supplementary Figure 8:** Culturing effects on MSC transcriptome. **(A)** Illustration of control groups examined. **(B)** Hallmark gene sets enrichment of LongNT vs. ShortNT. Results of GSEA Hallmark analysis show enriched gene sets (FDR q-value < 25% and p-val < 0.05). A positive normalized enrichment score (NES) values, marked in red, indicate enrichment in the LongNT phenotype. **(C)**. Circadian clock genes changed during normal cell growth (STRING network). Upregulated and downregulated following three days culturing are marked in red and blue accordingly.

## Tables and their legends

**Supplementary Table 1:**
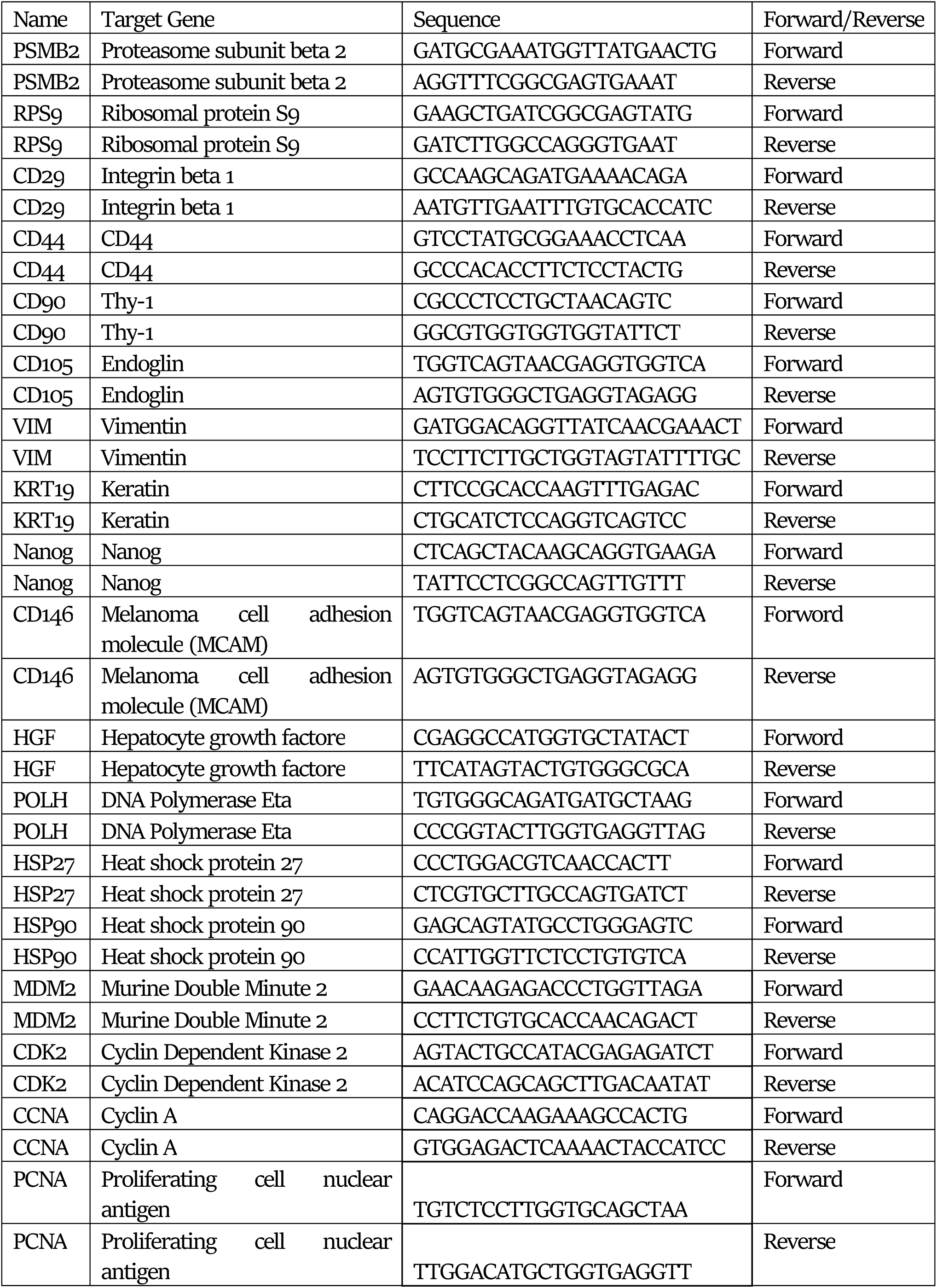
List of all bovine primers used in this study for gene expression analysis by RT-qPCR

**Supplementary Table 2:** Samples used for RNA-seq

| Sample Name | Total reads | Uniquely mapped reads (STAR) |
| --- | --- | --- |
| ShortNT-1 | 54,230,074 | 89.51% |
| ShortNT-2 | 39,577,082 | 90.10% |
| ShortNT-3 | 28,420,067 | 90.32% |
| LongNT-1 | 38,872,317 | 90.03% |
| LongNT-2 | 25,950,940 | 90.11% |
| ShortHS-1 | 48,638,628 | 89.98% |
| ShortHS-2 | 26,125,462 | 89.93% |
| ShortHS-3 | 28,842,747 | 89.88% |
| ShortHS_Recovery-1 | 45,307,646 | 90.08% |
| ShortHS_Recovery-2 | 20,743,593 | 89.81% |
| LongHS-1 | 39,755,614 | 90.03% |
| LongHS-2 | 30,679,249 | 89.87% |
| LongHS-3 | 26,106,742 | 90.52% |

**Supplementary Table 3:** Culturing conditions of the samples used in heat shock experiments

| Treatment name | Temperature (°C) | Time | Cell No. - Start | Cell No. - End |
| --- | --- | --- | --- | --- |
| ShortNT | 37°C | 6h | 250,000 | 329,212 |
| ShortHS | 40.5 °C | 6h | 250,000 | 320,365 |
| ShortHS_Recovery | 40.5 °C | 6h | 72,000 | 345,600 |
|  | 37 °C | 66h |  |  |
| LongNT | 37°C | 72h | 42,000 | 344,400 |
| LongHS | 40.5 °C | 72h | 72,000 | 104,982 |

**Supplementary Table S4:** Biological process upregulated in ShortHS - term convergence

**Supplementary Table 5:** List of GSEA GOBPs related to immune system and downregulated in LongHS

**Supplementary Table 6:** RNA-seq data analysis using DESeq

**Supplementary Table 7:** HS specific and common genes with no culturing effect

## Supplementary Materials and Methods

### RNA-seq data processing

Following sequencing, reads were trimmed for adaptors and poly A using Cutadapt, checked for quality by FastQC, mapped to genome (Bos_taurus.ARS-UCD1.2) and counted using STAR and HTSeq accordingly. Further quality control of alignment sequencing data was done using Integrative Genomics Viewer (IVG) and Qualimap application (Okonechnikov et al., 2016; Robinson et al., 2011). Normalization of the counts and differential expression analysis were performed using DESeq2 package in RStudio (RStudio Team, 2022). Total of 27,607 genes were sequenced. Genes with less then 5 counts in total per all treatments were removed, and we left with 16,595 genes for further downstream analyses (Supplementary Table S6). PCA, hierarchical and K-means clustering as well as other data visualizations, were performed using Rstudio.

### Functional enrichment analysis for immediate HS treatments (ShortHS & LongHS)

To get a comprehensive understanding of all the changes that occurred following the HS, we delved into the functional enrichment analysis using GSEA, g:Profiler and ClueGo tools, as recommended by Reimand et. al. (Bindea et al., 2009; Reimand et al., 2019; Zielniok et al., 2021).

Initially, genes with adjusted p-value (padj) < 0.05 and log2 Fold Change log2(FC) >1 were defined as DEGs. Gene Ontology (GO), Kyoto Encyclopedia of Genes and Genomes (KEGG) and Reactome pathways analysis was performed on DEGs using g:Profiler version e103_eg50_p15_eadf141 and ClueGO v2.5.7 (Cytoscape plug-in). In both tools default parameters with pv < 0.05 and bovine genome ARS-UCD1.2 were used. Analysis was performed for annotated genes only. In g:Profiler the analysis results were then filtered to display only terms sized between 5 and 350 genes, unless specified otherwise. DEG analysis was used in Figures 1-3 and supplementary figure S3.

Afterwards we continued the enrichment analysis using Gene Set Enrichment Analysis (GSEA) software. Since this is a human database, bovine gene names matching our gene id were taken from Ensemble BioMart ARS-UCD1.2. dataset and cross referenced with human orthologs from bioDBnet (biological DataBase network). Into GSEA, we entered the ranked list of all available genes from DESeq analysis (16,595) who have human ortholog (14,205). The enriched Hallmarks and GO pathways with a false discovery rate (FDR) less than 25% and p nominal value of less than were considered significantly meaningful. Settings used; 1000 permutations, collapse dataset to gene symbols - false, permutation type - gene set, default parameters in basic and advanced fields. GSEA analysis was used in Figures 2-3,5C and supplementary figures S7-8.

### Functional enrichment analysis for long term effects of HS

At the end, we wanted to check for any long-term effects of short HS and to compare between different HS groups. To this end, we re-analysed the data.

First, to eliminate culturing effect and focus on genes influenced by heat stress alone, the genes that changed between controls groups (Short and Long NT) were removed from the analysis. For this, all 16,595 sequenced genes were clustered, and 2 clusters (cluster 1 and 8 in Figure S9) were removed from gene list. Therefore 11679 genes left for further analysis. From 11,679 genes only 3,993 genes are with padj < 0.05 between ShortNT control and HS as shown in figure 4B. These 3,993 were used for functional enrichment analysis of long-term effect of HS (Figure 4), using g: Profiler e106_eg53_p16_65fcd97 with same parameters as stated previously.

**Figure S9:**
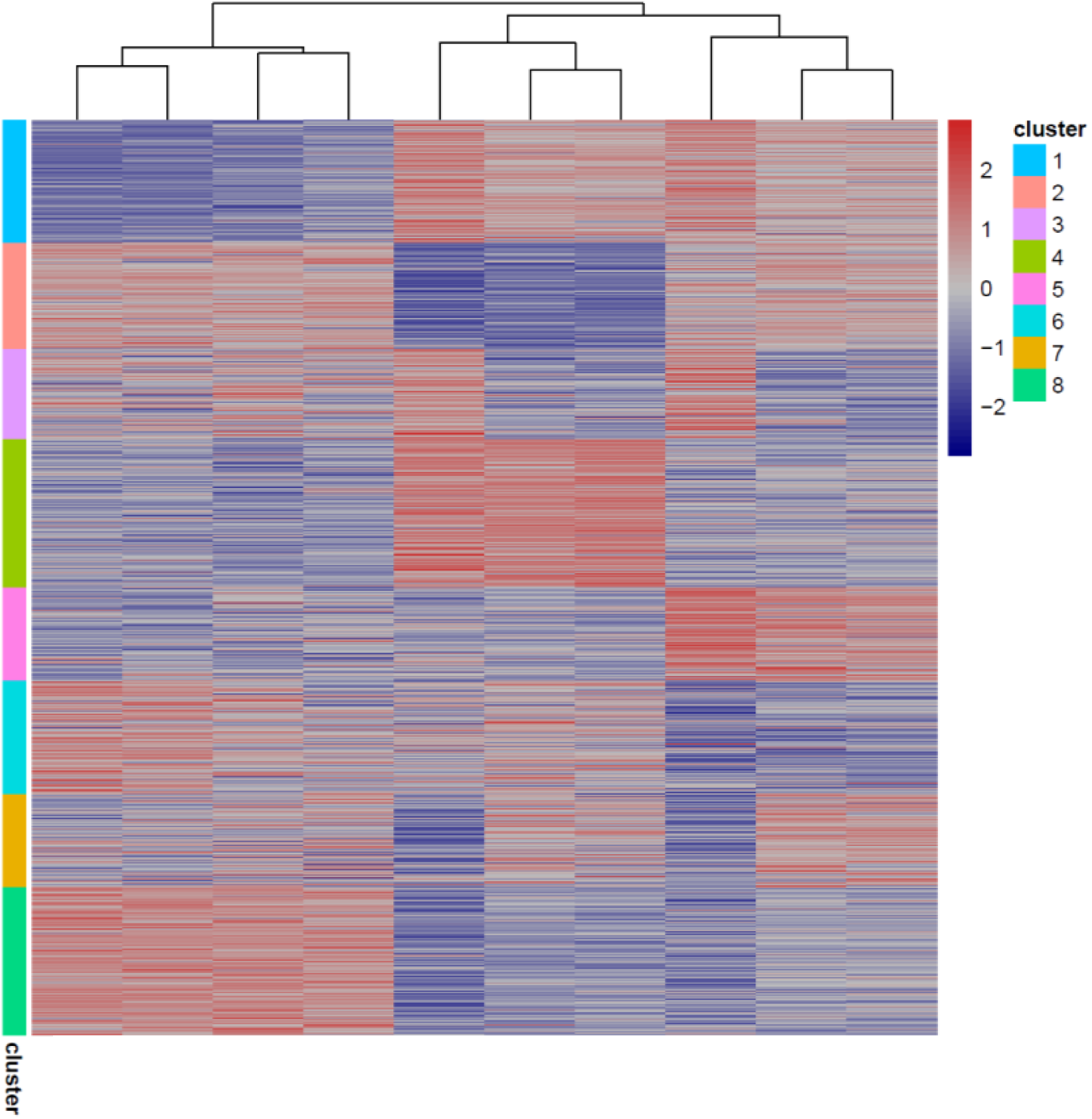
K-means clustering of 16595 genes from the sequencing. Cluster 1 and 8 were removed from gene list, Total of 11679 out of 16595 genes left for further analysis.

Since following ShortHS there was upregulation of many epigenetic changes (Figure 4F), we wanted to check if there’s any enrichment of specific epigenetic complexes. For this analysis was used complexes gene list from EpiFactors Database (Medvedeva et al., 2015) as well GO annotations of MGI database (Mouse Genome Informatics): MLL (GO:0044665, GO:0044666), HAT (GO:0000123) and HMT (GO:0035097). Bivalent genes enrichment was done by comparing our list to “3,868 bona-fide bivalent genes” in supplementary table No. 1 of (Mas et al., 2018). Since there’s no formal list of bivalent genes for any species, and there’s published diversity even within the same species, we chose to use Mas et al list of mouse bona fide bivalent since majority of them are also MLL2 targets. Enrichment calculation is explained in the statistical analysis section.

### Functional enrichment analysis for comparison between different HS groups

Comparison between different HS groups was done on 11,679 genes after removal of “culturing” genes as explained in previous section. Upregulated and downregulated genes with padj < 0.05 between each HS treatment and the relevant control, were intersected using Venn diagram to check for unique and common genes between the treatments (Figures 5A-B). List of genes with no culturing effect was prepared for each treatment (Supplementary Table S7), and enrichment analysis for specific pathways was done using The Graeber Lab online calculator for hypergeometric p value. Pathways analyzed: N00455: CDC25-Cell cycle G2/M (KEGG pathway), R-BTA-69002 – DNA Replication Pre-Initiation (Reactome), GO:0007049: Cell cycle (MGI database), GO:0060070: Canonical WNT signaling, GO 0048863: Stem cell differentiation, GO:0048468: Cell development, GO:0009888: Tissue development, GO:0007492: Endoderm development, GO:0007498: Mesoderm development, GO:0048762: Mesenchymal cell differentiation, GO:0002526: Acute Inflammatory response, GO:1905517: Macrophage migration, GO:0032635. Cytokine pathway list was taken from PathCards v5.7.551.0. This analysis was used in Figure 5. Enrichment calculation is explained in the statistical analysis section.

### Additional analysis

In order to mark the DEGs influenced by different heat shock treatments, KEGG pathway maps were downloaded from KEGG pathway database and marked with colors for down (blue) and up (red) regulated genes on each pathway (cell cycle - bta041100, adhesion molecules - bta04514, axon guidance - bta04360, DNA replication - bta03030, Oxidative phosphorylation - bta00190, Glycolysis - bta00010, Wnt signaling - bta04310). String datasets was used for functional protein association networks. SVG formats were further processed in Inkscape graphic software 1.1.2.

### Statistical Analysis

GraphPad Prism (La Jolla, CA, United States) was used for statistical analysis and visualization of RT-PCR. P < 0.05 was considered statistically significant. For Enrichment analysis hypergeometric p value was calculated using The Graeber Lab online calculator 2009 © (https://systems.crump.ucla.edu/hypergeometric/). Parameters used for hypergeometric test: N (population size) = 27607; M (number of successes in population) = number of genes in the tested pathway, (e.g., in MLL1-2 list contains total of 32 genes); s (sample size) = number of genes in specific treatment or cluster; k (number of successes) = number of genes present from pathway list in specific treatment or cluster.

### Tools and Database URLs

• IGV (https://software.broadinstitute.org/software/igv/)
• QualiMap (http://qualimap.conesalab.org/)
• RStudio Team (2020). RStudio: Integrated Development for R. RStudio, PBC, Boston, (https://www.rstudio.com/)
• Ensembl BioMart (https://www.ensembl.org/info/data/biomart/)
• bioDBnet (https://biodbnet-abcc.ncifcrf.gov/)
• GSEA (http://software.broadinstitute.org/gsea/)
• g:Profiler (https://biit.cs.ut.ee/gprofiler/)
• Cytoscape (http://www.cytoscape.org/) - ClueGO application
• EpiFactors Database (https://epifactors.autosome.org/)
• MGI database (http://www.informatics.jax.org/vocab/gene_ontology/)
• The Graeber Lab online calculator (https://systems.crump.ucla.edu/hypergeometric/)
• BioVenn (https://www.biovenn.nl/index.php)
• Inkscape (https://inkscape.org/)
• String database (https://string-db.org/)

### RNA-seq analysis workflow

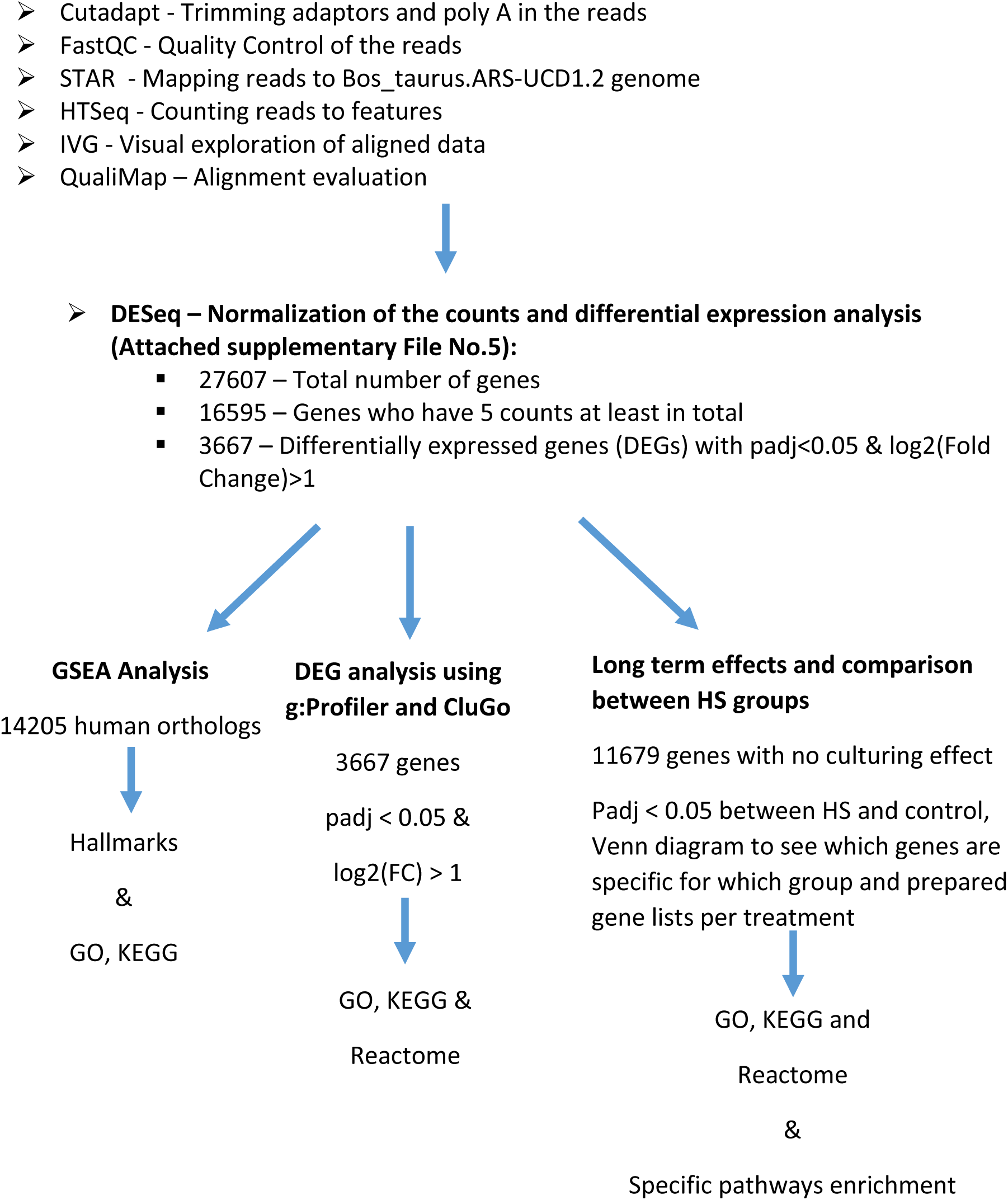

## References

Abdal Dayem, A., Lee, S., Y. Choi, H., & Cho, S. G. (2018). The Impact of Adhesion Molecules on the In Vitro Culture and Differentiation of Stem Cells. Biotechnology Journal, 13(2), 1700575. https://doi.org/10.1002/BIOT.201700575

Agarwal, A., Virk, G., Ong, C., & Plessis, S. S. du. (2014). Effect of Oxidative Stress on Male Reproduction. The World Journal of Men’s Health, 32(1), 1. https://doi.org/10.5534/WJMH.2014.32.1.1

Antonioli, E., Torres, N., Ferretti, M., de Azevedo Piccinato, C., & Sertie, A. L. (2019). Individual response to mTOR inhibition in delaying replicative senescence of mesenchymal stromal cells. PLoS ONE, 14(1). https://doi.org/10.1371/JOURNAL.PONE.0204784

Asseng, S., Spänkuch, D., Hernandez-Ochoa, I. M., & Laporta, J. (2021). The upper temperature thresholds of life. The Lancet Planetary Health, 5(6), e378–e385. https://doi.org/10.1016/S2542-5196(21)00079-6

Bagath, M., Krishnan, G., Devaraj, C., Rashamol, V. P., Pragna, P., Lees, A. M., & Sejian, V. (2019). The impact of heat stress on the immune system in dairy cattle: A review. Research in Veterinary Science, 126, 94–102. https://doi.org/10.1016/J.RVSC.2019.08.011

Chen, S., Wang, J., Peng, D., Li, G., Chen, J., & Gu, X. (2018). Exposure to heat-stress environment affects the physiology, circulation levels of cytokines, and microbiome in dairy cows. Scientific Reports 2018 *8*:1, 8(1), 1–11. https://doi.org/10.1038/s41598-018-32886-1

Choudhery, M. S. (2021). Strategies to improve regenerative potential of mesenchymal stem cells. World Journal of Stem Cells, 13(12), 1845. https://doi.org/10.4252/WJSC.V13.I12.1845

Collier, R. J., Stiening, C. M., Pollard, B. C., VanBaale, M. J., Baumgard, L. H., Gentry, P. C., & Coussens, P. M. (2006). Use of gene expression microarrays for evaluating environmental stress tolerance at the cellular level in cattle. Journal of Animal Science, 84(suppl_13), E1–E13. https://doi.org/10.2527/2006.8413_SUPPLE1X

Dado-Senn, B., Laporta, J., & Dahl, G. E. (2020). Carry over effects of late-gestational heat stress on dairy cattle progeny. Theriogenology, 154, 17–23. https://doi.org/10.1016/J.THERIOGENOLOGY.2020.05.012

Dalton, S., & Coverdell, P. D. (2015). Linking the cell cycle to cell fate decisions. Trends in Cell Biology, 25(10), 592. https://doi.org/10.1016/J.TCB.2015.07.007

de Barros, F. R. O., & Paula-Lopes, F. F. (2018). Cellular and epigenetic changes induced by heat stress in bovine preimplantation embryos. Molecular Reproduction and Development, 85(11), 810–820. https://doi.org/10.1002/MRD.23040

Dominici, M., Le Blanc, K., Mueller, I., Slaper-Cortenbach, I., Marini, F. C., Krause, D. S., Deans, R. J., Keating, A., Prockop, D. J., & Horwitz, E. M. (2006). Minimal criteria for defining multipotent mesenchymal stromal cells. The International Society for Cellular Therapy position statement. Cytotherapy, 8(4), 315–317. https://doi.org/10.1080/14653240600855905

Dou, J., Cánovas, A., Brito, L. F., Yu, Y., Schenkel, F. S., & Wang, Y. (2021). Comprehensive RNA-Seq Profiling Reveals Temporal and Tissue-Specific Changes in Gene Expression in Sprague– Dawley Rats as Response to Heat Stress Challenges. Frontiers in Genetics, 12, 420. https://doi.org/10.3389/FGENE.2021.651979/BIBTEX

Ermolaeva, M., Neri, F., Ori, A., & Rudolph, K. L. (2018). Cellular and epigenetic drivers of stem cell ageing. Nature Reviews Molecular Cell Biology 2018 19:9, 19(9), 594–610. https://doi.org/10.1038/s41580-018-0020-3

Fang, H., Kang, L., Abbas, Z., Hu, L., Chen, Y., Tan, X., Wang, Y., & Xu, Q. (2021). Identification of key Genes and Pathways Associated With Thermal Stress in Peripheral Blood Mononuclear Cells of Holstein Dairy Cattle. Frontiers in Genetics, 12, 910. https://doi.org/10.3389/FGENE.2021.662080/BIBTEX

Gao, S. T., Ma, L., Zhou, Z., Zhou, Z. K., Baumgard, L. H., Jiang, D., Bionaz, M., & Bu, D. P. (2019). Heat stress negatively affects the transcriptome related to overall metabolism and milk protein synthesis in mammary tissue of midlactating dairy cows. Physiological Genomics, 51(8), 400–409. https://doi.org/10.1152/PHYSIOLGENOMICS.00039.2019/ASSET/IMAGES/LARGE/ZH70091943690006.JPEG

Garner, J. B., Chamberlain, A. J., Vander Jagt, C., Nguyen, T. T. T., Mason, B. A., Marett, L. C., Leury, B. J., Wales, W. J., & Hayes, B. J. (2020). Gene expression of the heat stress response in bovine peripheral white blood cells and milk somatic cells in vivo. Scientific Reports 2022 *10*:1, *10*(1), 1–12. https://doi.org/10.1038/s41598-020-75438-2

Harikumar, A., & Meshorer, E. (2015). “Histones and Chromatin” Review series: Chromatin remodeling and bivalent histone modifications in embryonic stem cells. EMBO Reports, 16(12), 1609. https://doi.org/10.15252/EMBR.201541011

Hass, R., Kasper, C., Böhm, S., & Jacobs, R. (2011). Different populations and sources of human mesenchymal stem cells (MSC): A comparison of adult and neonatal tissue-derived MSC. Cell Communication and Signaling : CCS, 9(1), 12. https://doi.org/10.1186/1478-811X-9-12

Isik, B., Thaler, R., Goksu, B. B., Conley, S. M., Al-Khafaji, H., Mohan, A., Afarideh, M., Abumoawad, M., Zhu, X. Y., Krier, J. D., Saadiq, I. M., Tang, H., Eirin, A., Hickson, L. T. J., van Wijnen, A. J., Textor, S. C., Lerman, L. O., & Herrmann, S. M. (2021). Hypoxic preconditioning induces epigenetic changes and modifies swine mesenchymal stem cell angiogenesis and senescence in experimental atherosclerotic renal artery stenosis. Stem Cell Research and Therapy, 12(1), 1–13. https://doi.org/10.1186/S13287-021-02310-Z

Johnson, S. C., Rabinovitch, P. S., & Kaeberlein, M. (2013). mTOR is a key modulator of ageing and age-related disease. Nature, 493(7432), 338. https://doi.org/10.1038/NATURE11861

Kawano, K., Sakaguchi, K., Madalitso, C., Ninpetch, N., Kobayashi, S., Furukawa, E., Yanagawa, Y., & Katagiri, S. (2022). Effect of heat exposure on the growth and developmental competence of bovine oocytes derived from early antral follicles. Scientific Reports 2022 *12*:1, 12(1), 1–14. https://doi.org/10.1038/s41598-022-12785-2

Key, N., Sneeringer, S., & Marquardt, D. (2014). Climate Change, Heat Stress, and U.S. Dairy Production. SSRN Electronic Journal, 175. https://doi.org/10.2139/ssrn.2506668

Khalid, A. B., Pence, J., Suthon, S., Lin, J., Miranda-Carboni, G. A., & Krum, S. A. (2021). GATA4 regulates mesenchymal stem cells via direct transcriptional regulation of the WNT signalosome. Bone, 144, 115819. https://doi.org/10.1016/J.BONE.2020.115819

Khan, A., Dou, J., Wang, Y., Jiang, X., Khan, M. Z., Luo, H., Usman, T., & Zhu, H. (2020). Evaluation of heat stress effects on cellular and transcriptional adaptation of bovine granulosa cells. Journal of Animal Science and Biotechnology, 11(1). https://doi.org/10.1186/S40104-019-0408-8

Kitagawa, Y., Suzuki, K., Yoneda, A., & Watanabe, T. (2004). Effects of oxygen concentration and antioxidants on the in vitro developmental ability, production of reactive oxygen species (ROS), and DNA fragmentation in porcine embryos. Theriogenology, 62(7), 1186–1197. https://doi.org/10.1016/J.THERIOGENOLOGY.2004.01.011

Kou, M., Huang, L., Yang, J., Chiang, Z., Chen, S., Liu, J., Guo, L., Zhang, X., Zhou, X., Xu, X., Yan, X., Wang, Y., Zhang, J., Xu, A., Tse, H., & Lian, Q. (2022). Mesenchymal stem cell-derived extracellular vesicles for immunomodulation and regeneration: a next generation therapeutic tool? Cell Death & Disease 2022 13:7, 13(7), 1–16. https://doi.org/10.1038/s41419-022-05034-x

Kuroda, Y., & Dezawa, M. (2014). Mesenchymal stem cells and their subpopulation, pluripotent muse cells, in basic research and regenerative medicine. Anatomical Record, 297(1), 98–110. https://doi.org/10.1002/ar.22798

Li, J., Labbadia, J., & Morimoto, R. I. (2017). Rethinking HSF1 in Stress, Development, and Organismal Health. Trends in Cell Biology, 27(12), 895–905. https://doi.org/10.1016/J.TCB.2017.08.002

Madrigal, M., Rao, K. S., & Riordan, N. H. (2014). A review of therapeutic effects of mesenchymal stem cell secretions and induction of secretory modification by different culture methods. Journal of Translational Medicine, 12(1), 1–14. https://doi.org/10.1186/s12967-014-0260-8

Mas, G., Blanco, E., Ballaré, C., Sansó, M., Spill, Y. G., Hu, D., Aoi, Y., Le Dily, F., Shilatifard, A., Marti-Renom, M. A., & Di Croce, L. (2018). Promoter bivalency favors an open chromatin architecture in embryonic stem cells. Nature Genetics 2018 50:10, 50(10), 1452–1462. https://doi.org/10.1038/s41588-018-0218-5

Medvedeva, Y. A., Lennartsson, A., Ehsani, R., Kulakovskiy, I. V., Vorontsov, I. E., Panahandeh, P., Khimulya, G., Kasukawa, T., & Drabløs, F. (2015). EpiFactors: a comprehensive database of human epigenetic factors andcomplexes. Database: The Journal of Biological Databases and Curation, 2015. https://doi.org/10.1093/DATABASE/BAV067

Moise, S., Byrne, J. M., El Haj, A. J., & Telling, N. D. (2018). The potential of magnetic hyperthermia for triggering the differentiation of cancer cells. Nanoscale, 10(44), 20519–20525. https://doi.org/10.1039/C8NR05946B

Naaldijk, Y., Johnson, A. A., Ishak, S., Meisel, H. J., Hohaus, C., & Stolzing, A. (2015). Migrational changes of mesenchymal stem cells in response to cytokines, growth factors, hypoxia, and aging. Experimental Cell Research, 338(1), 97–104. https://doi.org/10.1016/J.YEXCR.2015.08.019

Nir & Ribarski-Chorev, D., Ribarski-Chorev, I., Shimoni, C., Strauss, C., Frank, J., & Schlesinger, S. (2022). Antioxidants Attenuate Heat Shock Induced Premature Senescence of Bovine Mesenchymal Stem Cells. International Journal of Molecular Sciences, 23(10), 5750. https://doi.org/10.3390/ijms23105750

Noer, A., Lindeman, L. C., & Collas, P. (2009). Histone H3 modifications associated with differentiation and long-term culture of mesenchymal adipose stem cells. Stem Cells and Development, 18(5), 725–736. https://doi.org/10.1089/scd.2008.0189

Nowakowski, A., Drela, K., Rozycka, J., Janowski, M., & Lukomska, B. (2016). Engineered Mesenchymal Stem Cells as an Anti-Cancer Trojan Horse. Stem Cells and Development, 25(20), 1513–1531. https://doi.org/10.1089/scd.2016.0120

Olde Riekerink, R. G. M., Barkema, H. W., & Stryhn, H. (2007). The Effect of Season on Somatic Cell Count and the Incidence of Clinical Mastitis. Journal of Dairy Science, 90(4), 1704–1715. https://doi.org/10.3168/JDS.2006-567

Peters, A., Nawrot, T. S., & Baccarelli, A. A. (2021). Hallmarks of environmental insults. Cell, 184(6), 1455–1468. https://doi.org/10.1016/J.CELL.2021.01.043

Phinney, D. G., & Pittenger, M. F. (2017). Concise Review: MSC-Derived Exosomes for Cell-Free Therapy. STEMCELLS, 35(4), 851–858. https://doi.org/10.1002/STEM.2575

Pittenger, M. F., Mackay, A. M., Beck, S. C., Jaiswal, R. K., Douglas, R., Mosca, J. D., Moorman, M. A., Simonetti, D. W., Craig, S., & Marshak, D. R. (1999). Multilineage potential of adult human mesenchymal stem cells. Science, 284(5411), 143–147. https://doi.org/10.1126/SCIENCE.284.5411.143/SUPPL_FILE/983855S5_THUMB.GIF

Richter, K., Haslbeck, M., & Buchner, J. (2010). The Heat Shock Response: Life on the Verge of Death. Molecular Cell, 40(2), 253–266. https://doi.org/10.1016/J.MOLCEL.2010.10.006/ATTACHMENT/BFF2FC30-41D1-4EE0-9FE1-1D8B605F6F20/MMC1.PDF

Roth, Z. (2017). Effect of Heat Stress on Reproduction in Dairy Cows: Insights into the Cellular and Molecular Responses of the Oocyte. Http://Dx.Doi.Org/10.1146/Annurev-Animal-022516-022849, 5, 151–170. https://doi.org/10.1146/ANNUREV-ANIMAL-022516-022849

Roth, Z. (2020). Reproductive physiology and endocrinology responses of cows exposed to environmental heat stress - Experiences from the past and lessons for the present. Theriogenology, 155, 150–156. https://doi.org/10.1016/J.THERIOGENOLOGY.2020.05.040

Sammad, A., Luo, H., Hu, L., Zhu, H., & Wang, Y. (2022). Transcriptome Reveals Granulosa Cells Coping through Redox, Inflammatory and Metabolic Mechanisms under Acute Heat Stress. Cells, 11(9). https://doi.org/10.3390/CELLS11091443/S1

Saxton, R. A., & Sabatini, D. M. (2017). mTOR Signaling in Growth, Metabolism, and Disease. Cell, 168(6), 960–976. https://doi.org/10.1016/J.CELL.2017.02.004

Schultz, M. B., & Sinclair, D. A. (2016). When stem cells grow old: phenotypes and mechanisms of stem cell aging. Development, 143(1), 3–14. https://doi.org/10.1242/DEV.130633

Sharma, S., & Bhonde, R. (2020). Genetic and epigenetic stability of stem cells: Epigenetic modifiers modulate the fate of mesenchymal stem cells. Genomics, 112(5), 3615–3623. https://doi.org/10.1016/J.YGENO.2020.04.022

Shimoni, C., Goldstein, M., Ribarski-Chorev, I., Schauten, I., Nir, D., Strauss, C., & Schlesinger, S. (2020). Heat Shock Alters Mesenchymal Stem Cell Identity and Induces Premature Senescence. Frontiers in Cell and Developmental Biology, 8(September), 1–15. https://doi.org/10.3389/fcell.2020.565970

Stopp, S., Bornhäuser, M., Ugarte, F., Wobus, M., Kuhn, M., Brenner, S., & Thieme, S. (2013). Expression of the melanoma cell adhesion molecule in human mesenchymal stromal cells regulates proliferation, differentiation, and maintenance of hematopoietic stem and progenitor cells. Haematologica, 98(4), 505–513. https://doi.org/10.3324/HAEMATOL.2012.065201

Su, K. H., & Dai, C. (2017). mTORC1 senses stresses: Coupling stress to proteostasis. *BioEssays : News and Reviews in Molecular*, Cellular and Developmental Biology, 39(5). https://doi.org/10.1002/BIES.201600268

Toupadakis, C. A., Wong, A., Genetos, D. C., Cheung, W. K., Borjesson, D. L., Ferraro, G. L., Galuppo, L. D., Leach, J. K., Owens, S. D., & Yellowley, C. E. (2010). Comparison of the osteogenic potential of equine mesenchymal stem cells from bone marrow, adipose tissue, umbilical cord blood, and umbilical cord tissue. American Journal of Veterinary Research, 71(10), 1237–1245. https://doi.org/10.2460/AJVR.71.10.1237

Uccelli, A., & al., et. (2008). Mesenchymal stem cells in health and disease. Nat. Rev. Immunol., 8, 726–736.

Uccelli, A., Moretta, L., & Pistoia, V. (2008). Mesenchymal stem cells in health and disease. Nat Rev Immunol, 8(9), 726–736. https://doi.org/10.1038/nri2395

van Velthoven, C. T. J., & Rando, T. A. (2019). Stem Cell Quiescence: Dynamism, Restraint, and Cellular Idling. Cell Stem Cell, 24(2), 213. https://doi.org/10.1016/J.STEM.2019.01.001

Xue, Y., & Acar, M. (2018). Mechanisms for the epigenetic inheritance of stress response in single cells. Current Genetics, 64(6), 1221. https://doi.org/10.1007/S00294-018-0849-1

Yue, S., Wang, Z., Wang, L., Peng, Q., & Xue, B. (2020). Transcriptome Functional Analysis of Mammary Gland of Cows in Heat Stress and Thermoneutral Condition. Animals : An Open Access Journal from MDPI, 10(6), 1–18. https://doi.org/10.3390/ANI10061015

Zhang, Y., Ling, L., Ajay D/O Ajayakumar, A., Eio, Y. M., van Wijnen, A. J., Nurcombe, V., & Cool, S. M. (2022). FGFR2 accommodates osteogenic cell fate determination in human mesenchymal stem cells. Gene, 818, 146199. https://doi.org/10.1016/J.GENE.2022.146199

