## Supplemental Figures for "Short heat shock has a long-term effect on mesenchymal stem cells’ transcriptome"

### Supplementary Figure S1

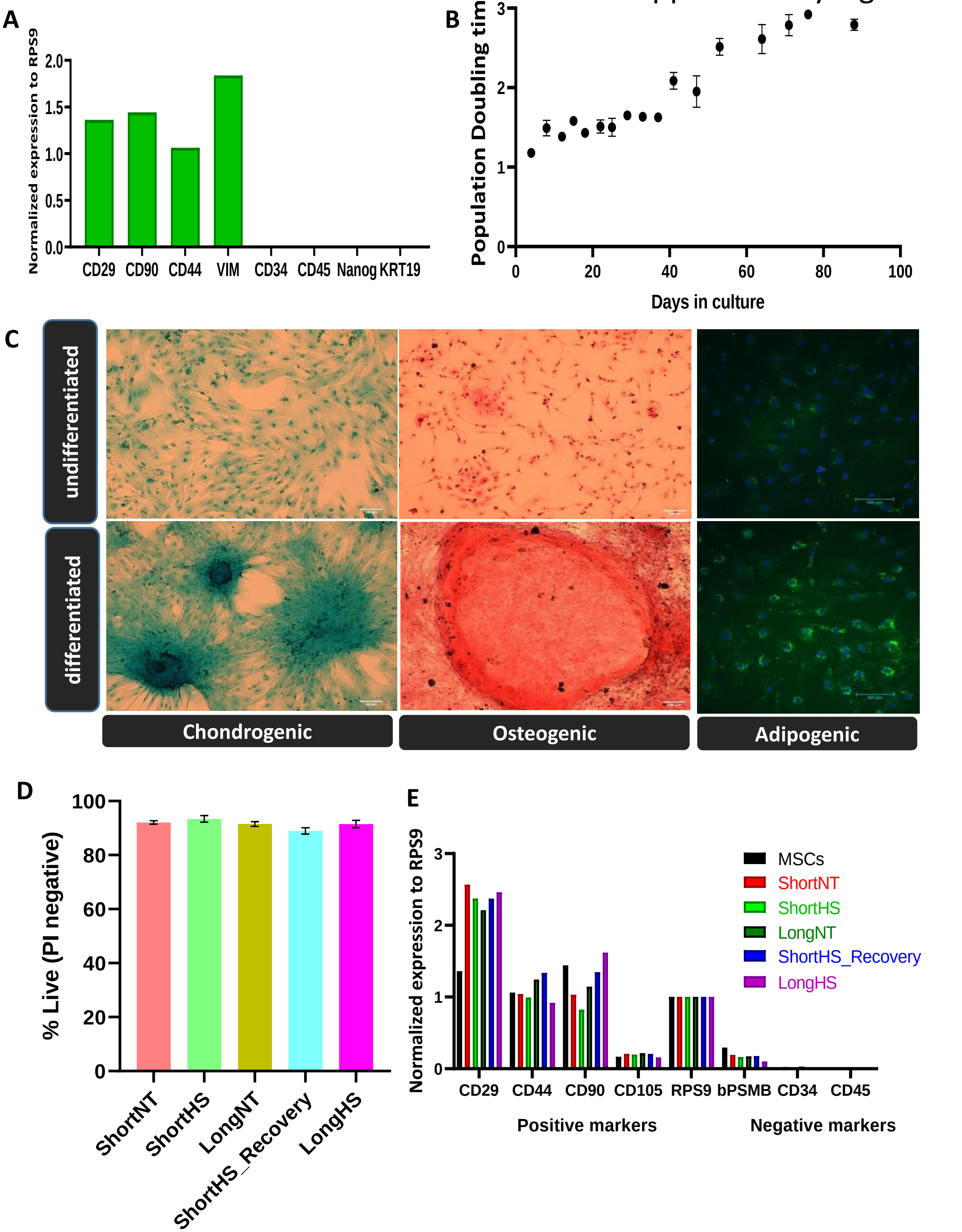

### Supplementary Figure S2

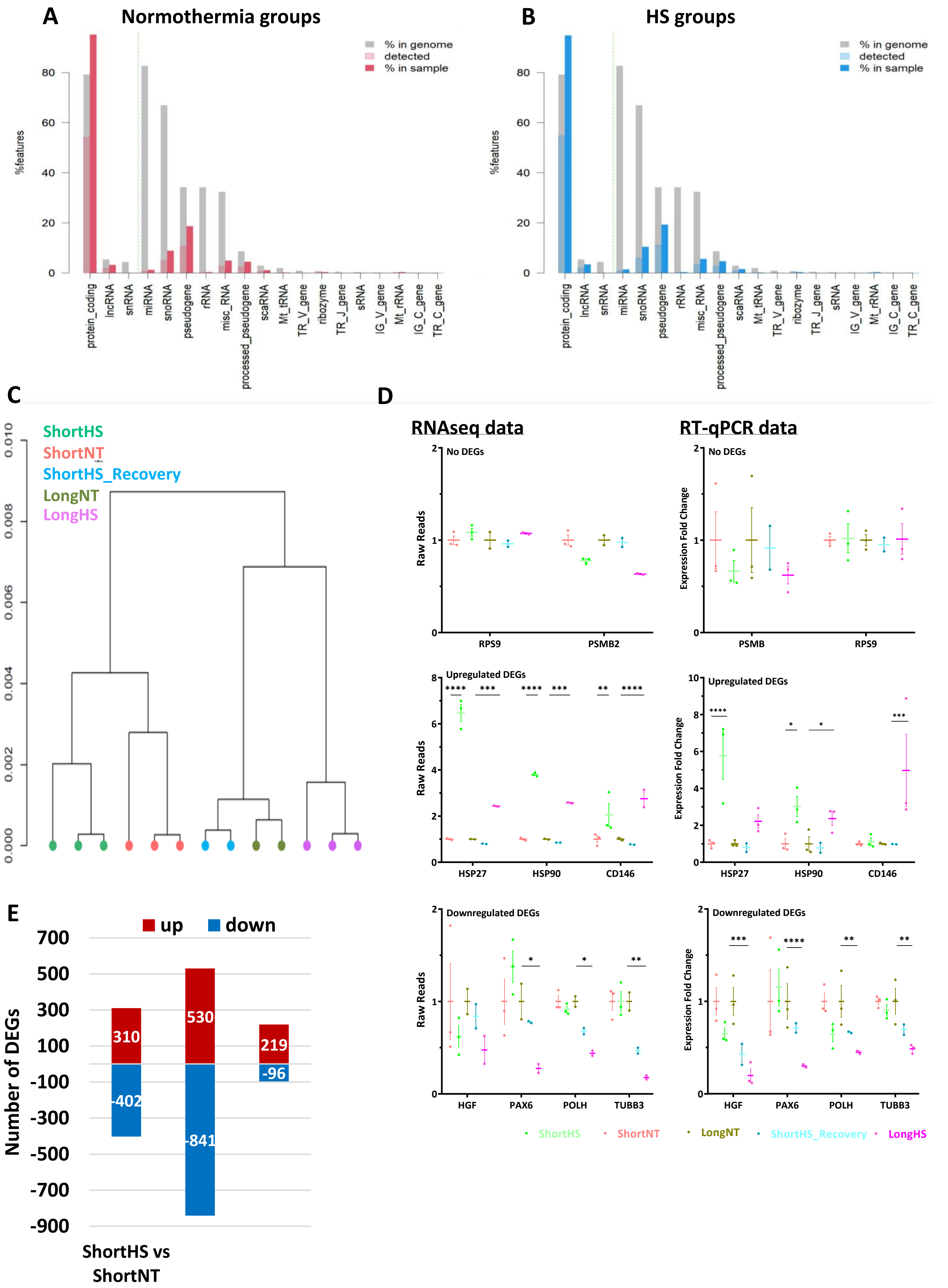

### Supplementary Figure S3

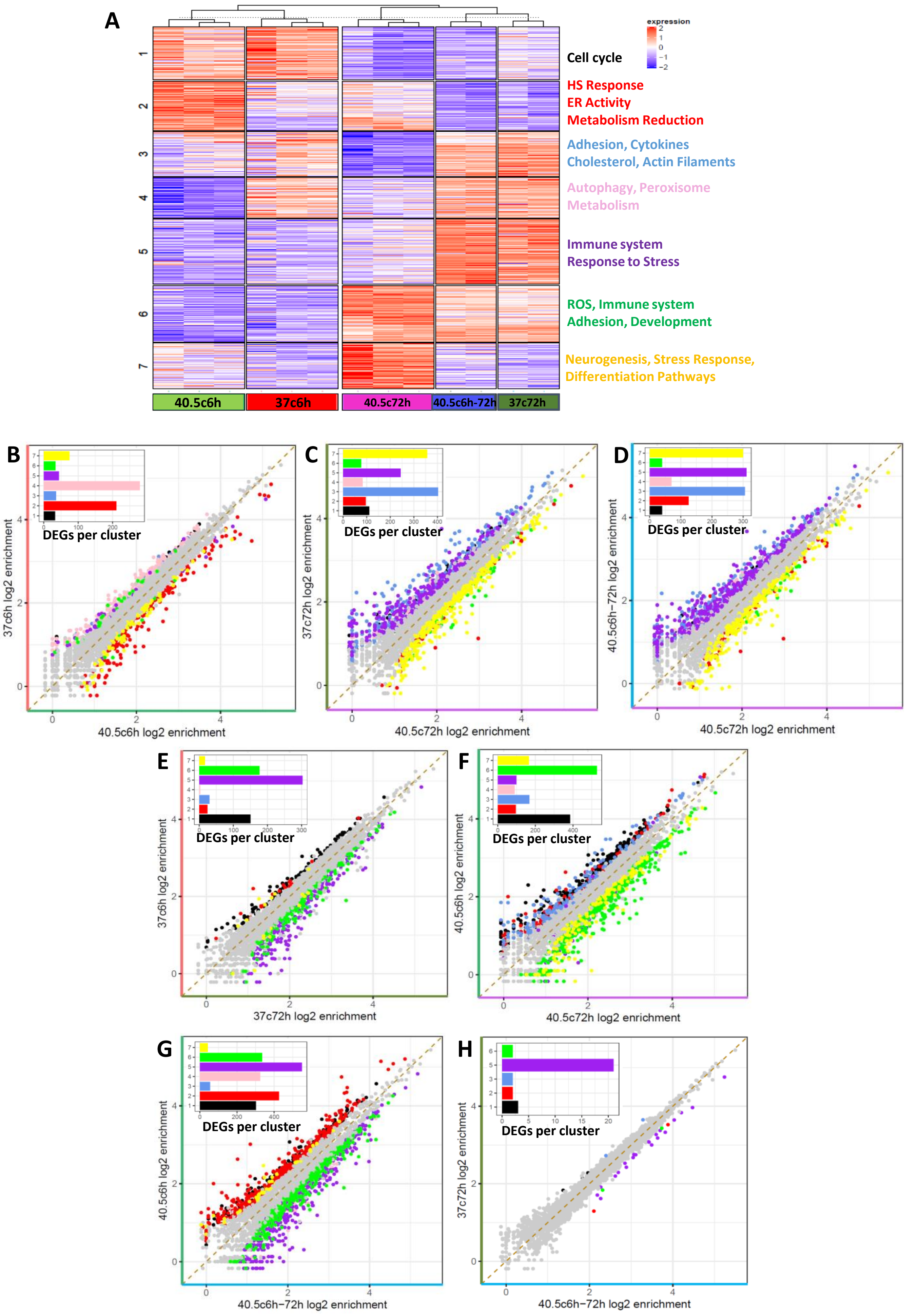

Supplementary Figure S4

#### A. Cell Cycle

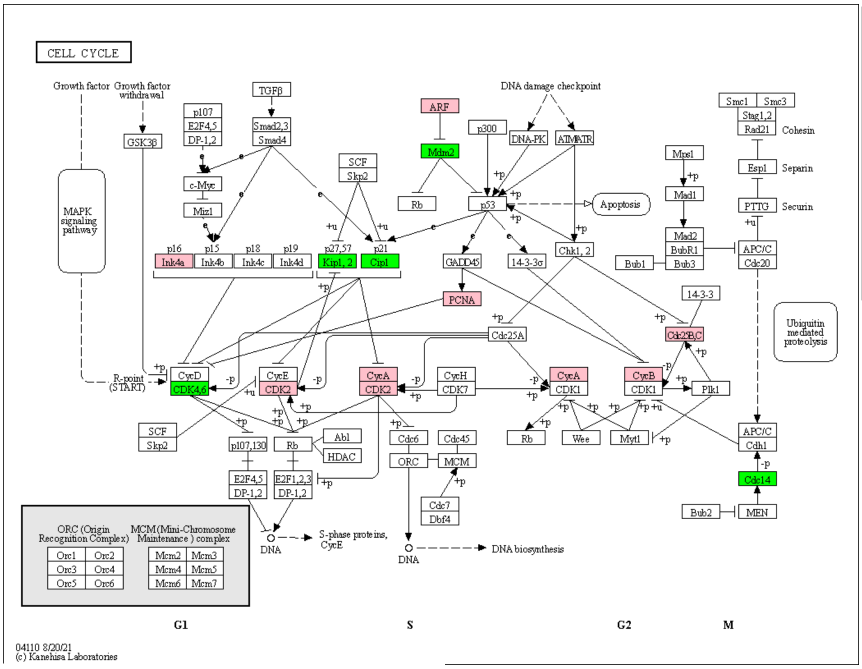


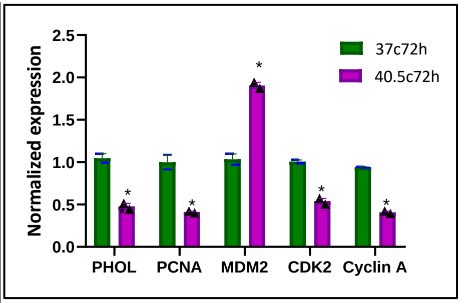


#### B. Cell Adhesion Molecules

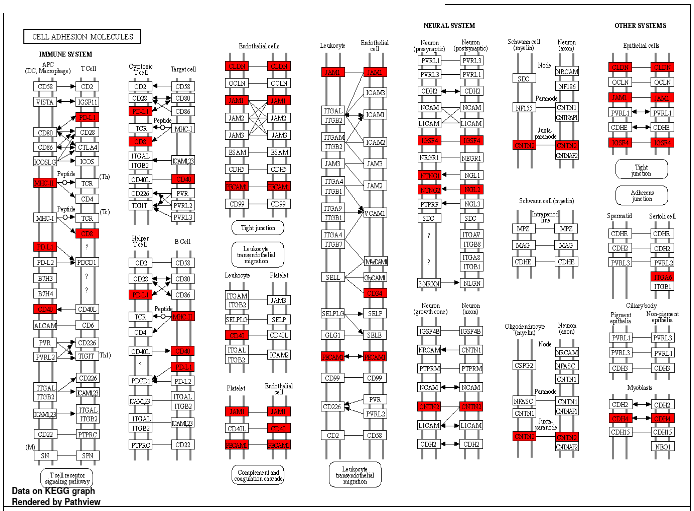


#### C. Axon Guidance

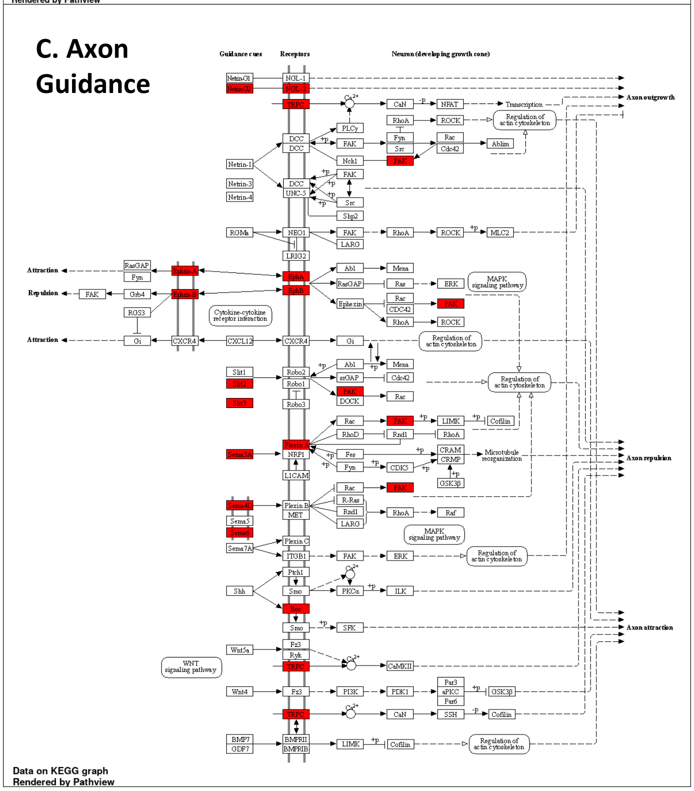

Supplementary Figure S5

**A**

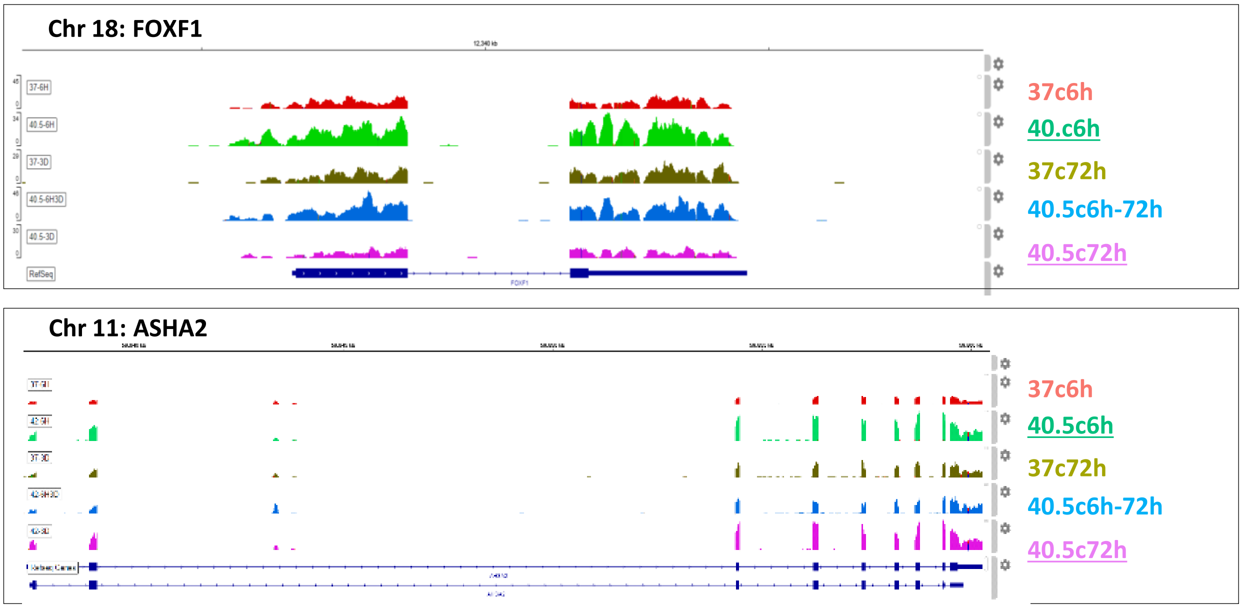


**B**

#### DNA Replication

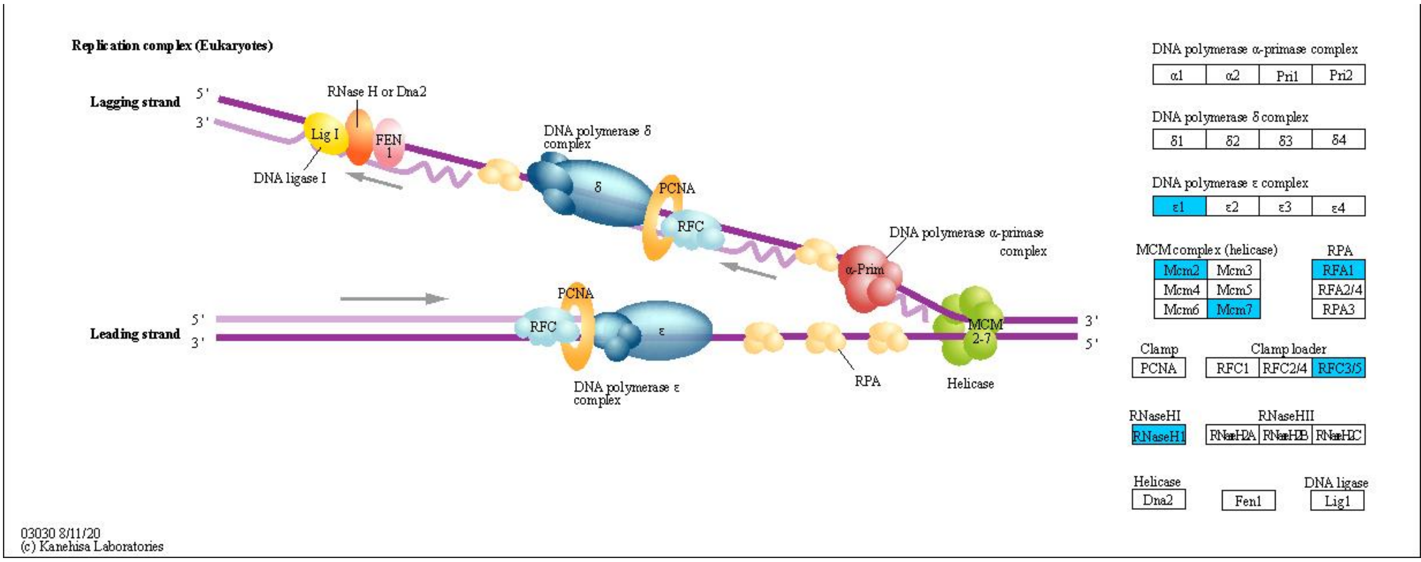

### Supplementary Figure S6

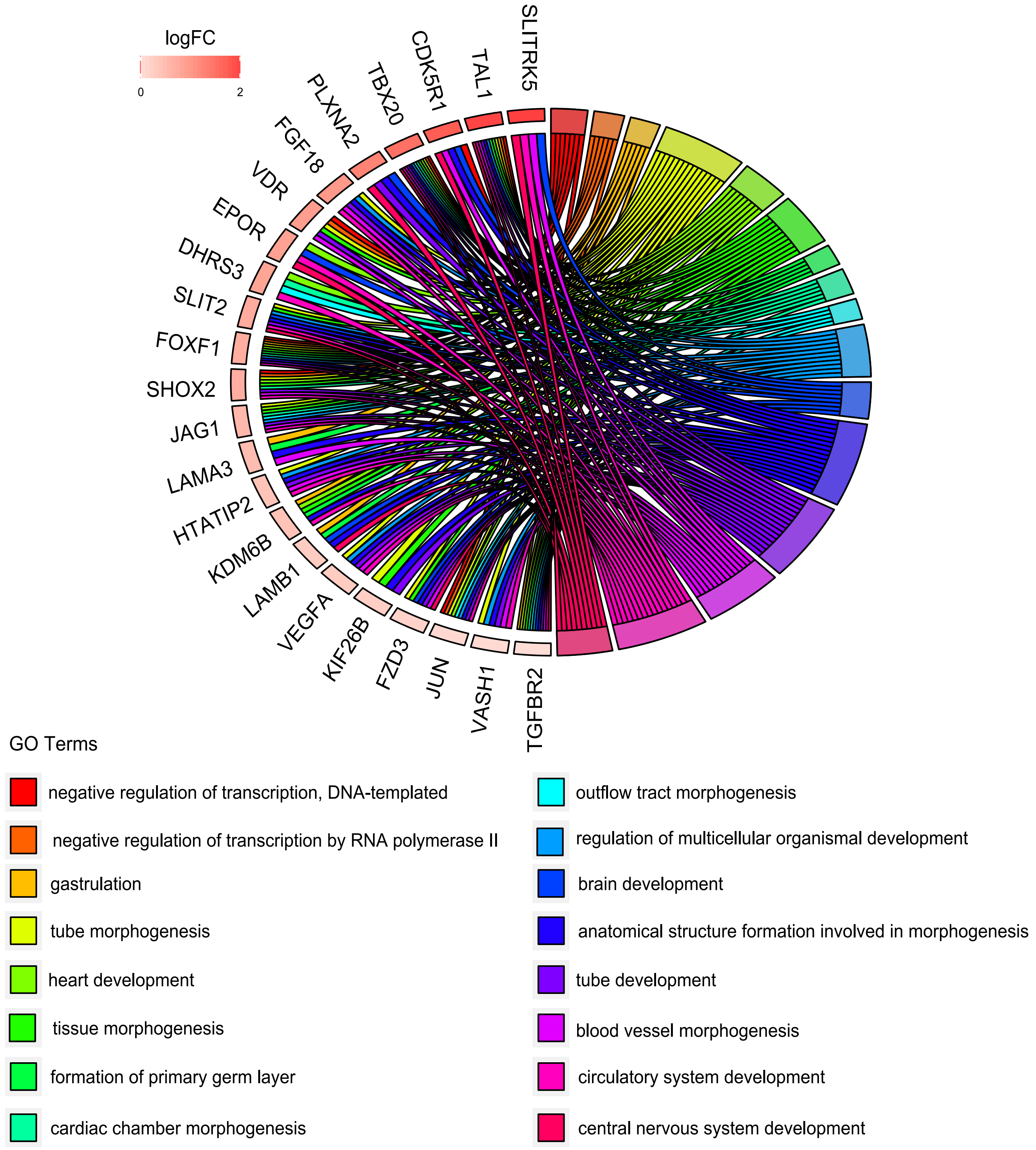

Supplementary Figure S7

A

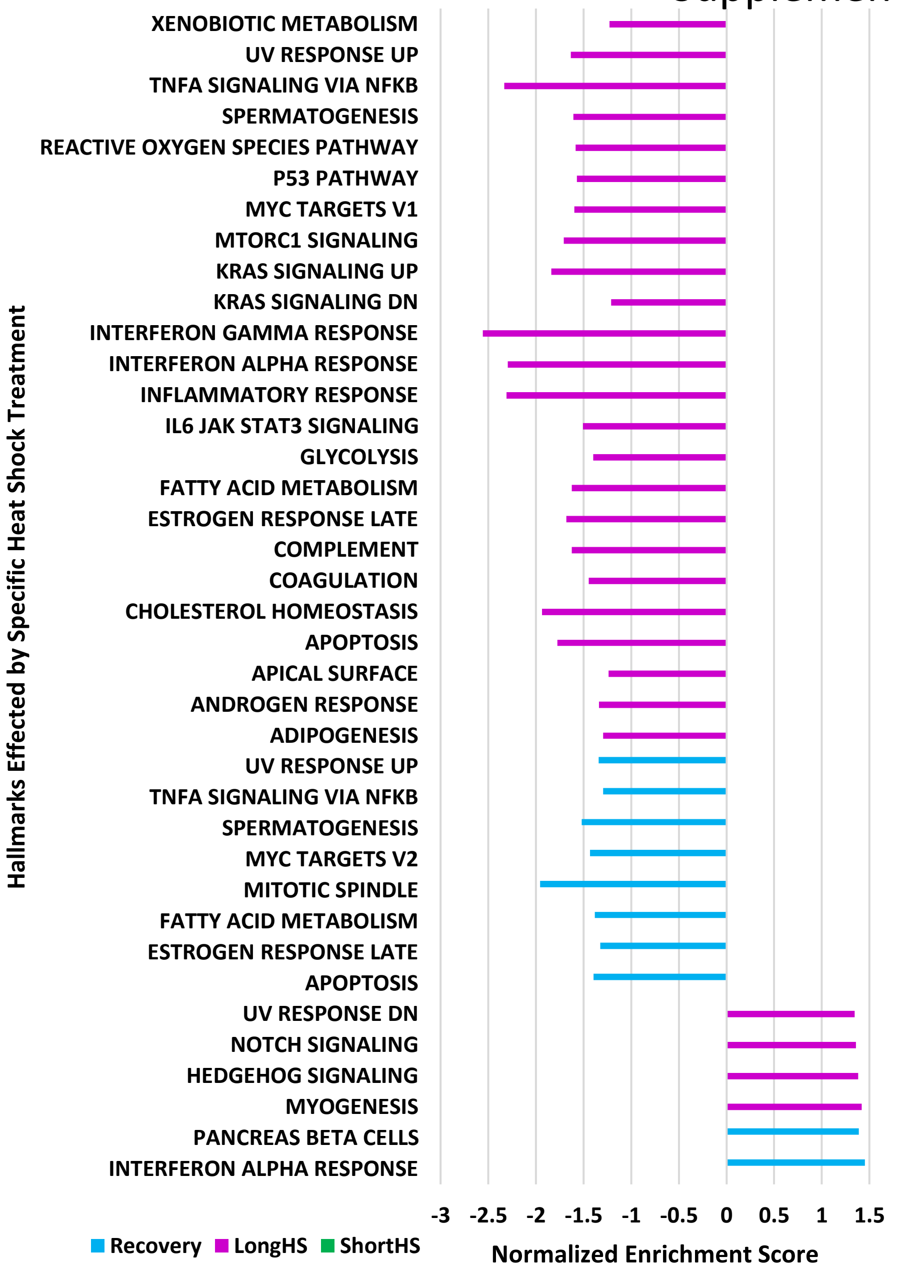


B

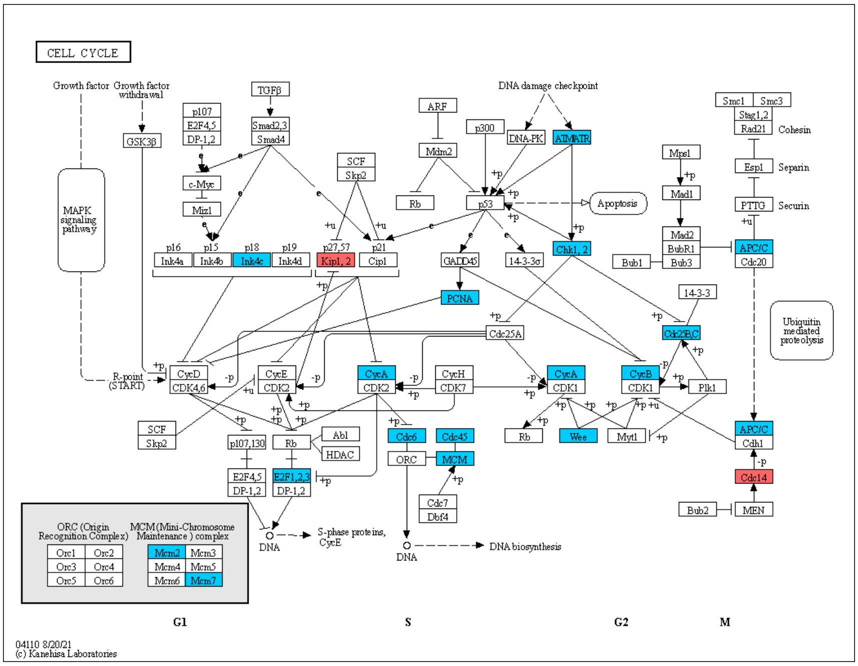

Supplementary Figure S7

C

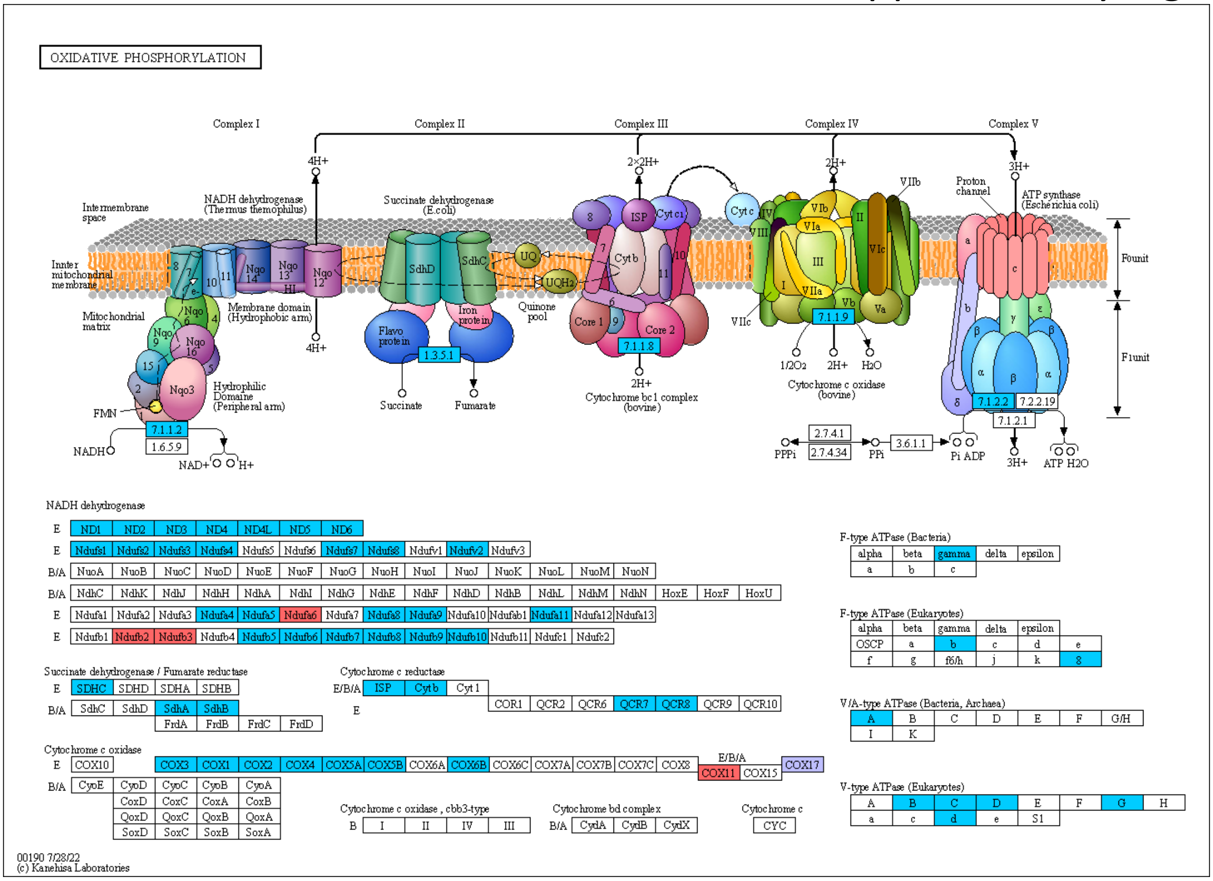


D

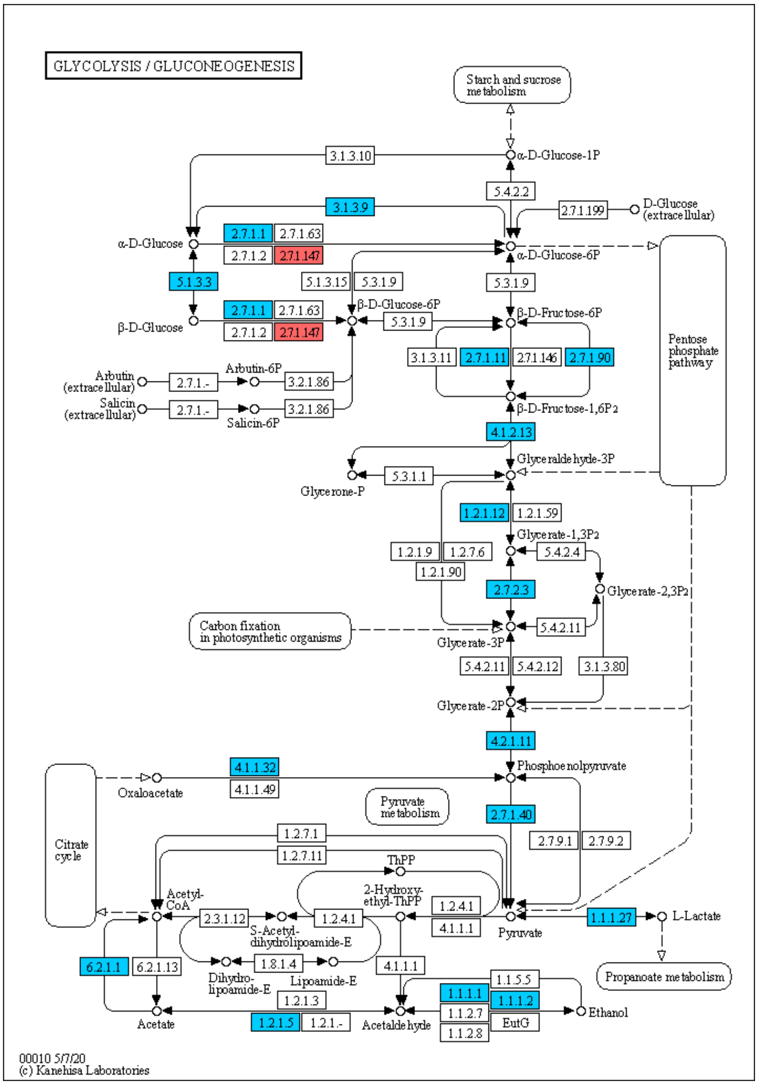

Supplementary Figure S7

E

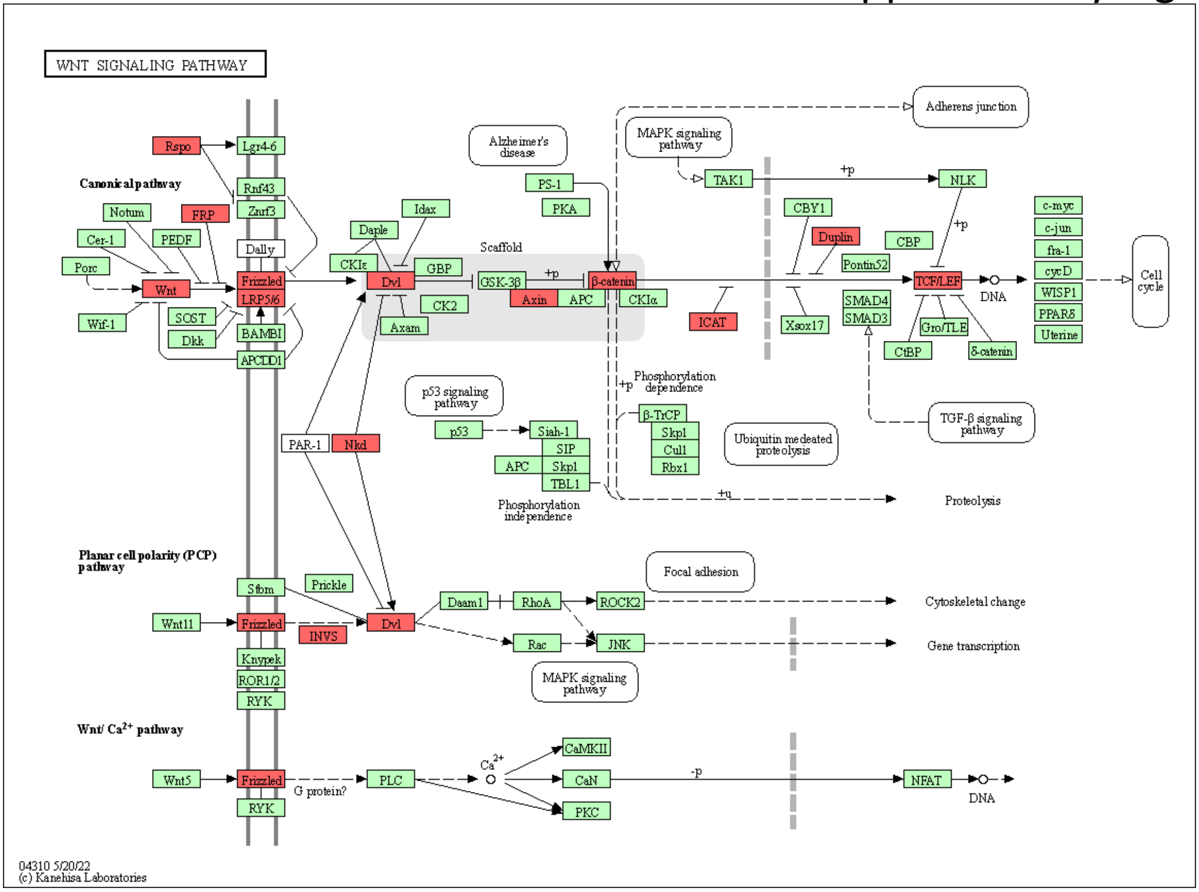


F

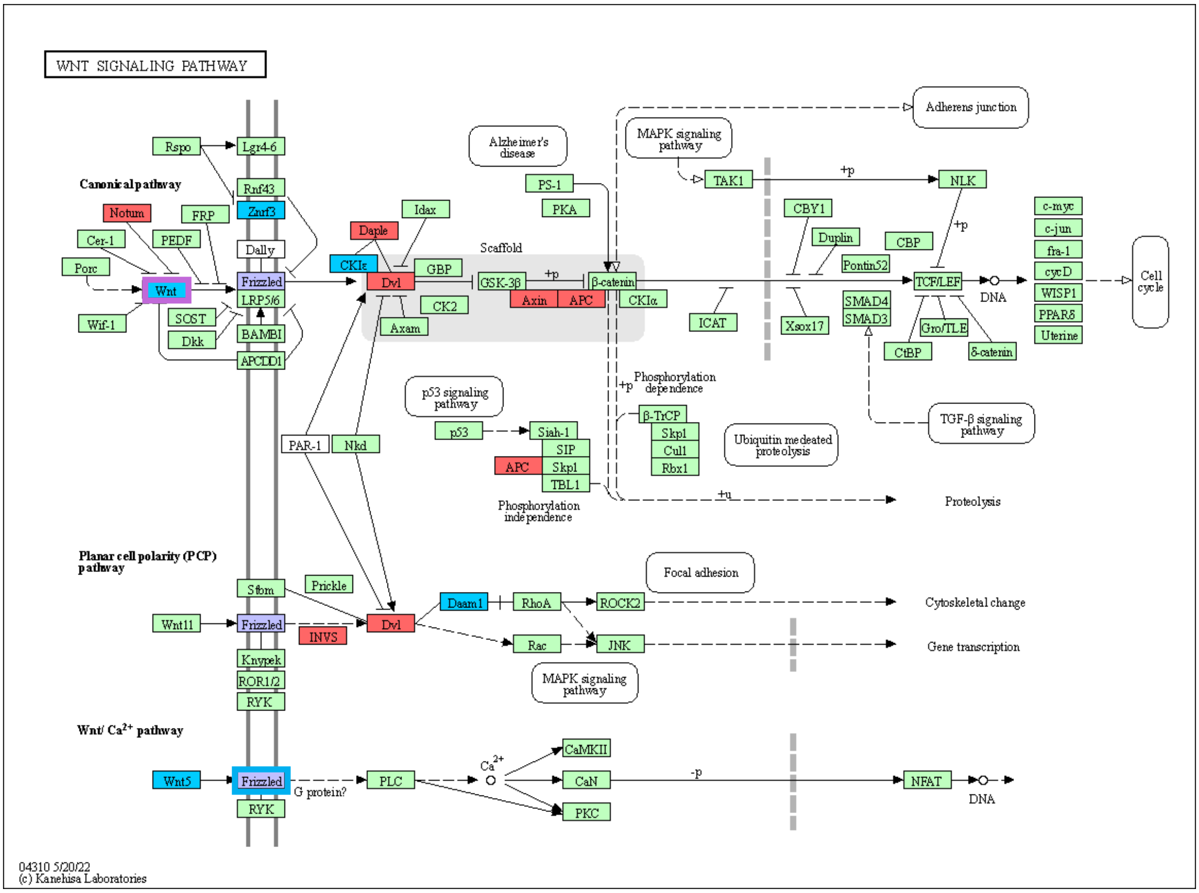

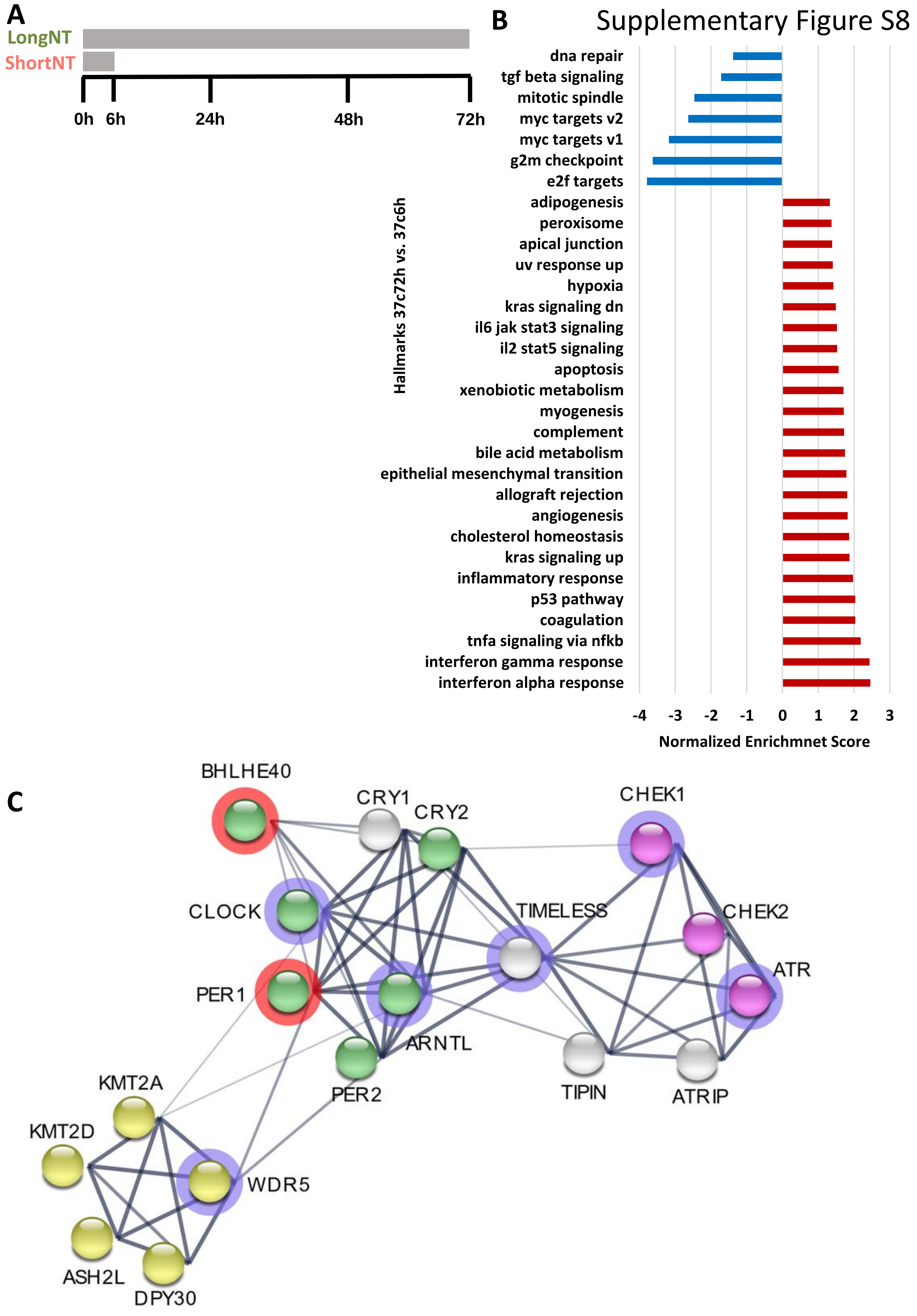
Supplementary Figure S8
A
LongNT
ShortNT
0h
6h
24h
48h
72h
B
Hallmarks 37c72h vs. 37c6h
dna repair
tgf beta signaling
mitotic spindle
myc targets v2
myc targets v1
g2m checkpoint
e2f targets
adipogenesis
peroxisome
apical junction
uv response up
hypoxia
kras signaling dn
il6 jak stat3 signaling
il2 stat5 signaling
apoptosis
xenobiotic metabolism
myogenesis
complement
bile acid metabolism
epithelial mesenchymal transition
allograft rejection
angiogenesis
cholesterol homeostasis
kras signaling up
inflammatory response
p53 pathway
coagulation
tnfa signaling via nfkb
interferon gamma response
interferon alpha response
-4
-3
-2
-1
0
1
2
3
Normalized Enrichmnet Score
C
BHLHE40
CRY1
CRY2
CHEK1
CLOCK
TIMELESS
CHEK2
ATR
PER1
ARNTL
PER2
TIPIN
ATRIP
KMT2A
KMT2D
WDR5
ASH2L
DPY30
